# Cancer-associated DNA Hypermethylation of Polycomb Targets Requires DNMT3A Dual Recognition of Histone H2AK119 Ubiquitination and the Nucleosome Acidic Patch

**DOI:** 10.1101/2024.03.18.585588

**Authors:** Kristjan H. Gretarsson, Stephen Abini-Agbomson, Susan L Gloor, Daniel N Weinberg, Jamie L McCuiston, Vishnu Udayakumar Sunitha Kumary, Allison R Hickman, Varun Sahu, Rachel Lee, Xinjing Xu, Natalie Lipieta, Samuel Flashner, Oluwatobi A. Adeleke, Irina K Popova, Hailey F Taylor, Kelsey Noll, Carolina Lin Windham, Danielle N Maryanski, Bryan J Venters, Hiroshi Nakagawa, Michael-Christopher Keogh, Karim-Jean Armache, Chao Lu

**Author notes:** Authors made equal contributions (Kristjan H. Gretarsson, Stephen Abini-Agbomson). Co-correspondence (M.C.K.); (K.-J.A.); (C.L.).

## Abstract

During tumor development, promoter CpG islands (CGIs) that are normally silenced by Polycomb repressive complexes (PRCs) become DNA hypermethylated. The molecular mechanism by which *de novo* DNA methyltransferase(s) catalyze CpG methylation at PRC-regulated regions remains unclear. Here we report a cryo-EM structure of the DNMT3A long isoform (DNMT3A1) N-terminal region in complex with a nucleosome carrying PRC1-mediated histone H2A lysine 119 monoubiquitination (H2AK119Ub). We identify regions within the DNMT3A1 N-terminus that bind H2AK119Ub and the nucleosome acidic patch. This bidentate interaction is required for effective DNMT3A1 engagement with H2AK119Ub-modified chromatin in cells. Furthermore, aberrant redistribution of DNMT3A1 to Polycomb target genes inhibits their transcriptional activation during cell differentiation and recapitulates the cancer-associated DNA hypermethylation signature. This effect is rescued by disruption of the DNMT3A1-acidic patch interaction. Together, our analyses reveal a binding interface critical for countering promoter CGI DNA hypermethylation, a major molecular hallmark of cancer.

## Introduction

An aberrant DNA methylation (DNAme; primarily 5-methylcytosine) landscape is a well-established pan-cancer molecular hallmark (1). As an example, megabase domains that are gene-poor, late-replicating and of low CG density gradually lose DNAme in tumors. Recent evidence suggests this progressive hypomethylation is coupled to cancer cell mitotic divisions and represents a byproduct of replicative aging (2). On the other hand, CpG islands (CGIs) – regions of high CG density found at >60% of human gene promoters (3) – are normally free of DNAme, yet become hypermethylated in cancer cells (4–6). Promoter CGI hypermethylation can lead to the silencing of downstream genes including many tumor suppressors such as *CDKN2A* and *BRCA1*, thereby critically contributing to cancer initiation and development (7–10).

The molecular mechanisms underlying cancer-associated promoter CGI hypermethylation remain a major focus in the field. Notably, multiple independent analyses of large cohorts of patient tumor samples demonstrate that promoter CGIs that gain DNAme are marked by tri-methylation of histone H3K27 (H3K27me3) in embryonic or tissue stem/progenitor cells (11–14). This histone post-translational modification (PTM) is established by the Polycomb Repressive Complex 2 (PRC2), which cooperates with another Polycomb complex (PRC1) and its enzymatic product H2AK119 monoubiquitination (H2AK119Ub) to maintain temporal silencing of genes involved in cell differentiation (15). Why Polycomb target genes are particularly vulnerable to DNA hypermethylation is not well understood. Previous studies have reported the genomic co-localization of Polycomb complex members and various DNA methyltransferases (DNMTs) (16,17). However, the molecular and structural bases for their interactions, if any, are unclear.

DNMT3A is one of the two *de novo* DNMTs in mammalian cells (18). *DNMT3A* encodes two major isoforms (19), with short DNMT3A2 specifically expressed in early embryonic development, while long DNMT3A1 is universally expressed across somatic tissues. The catalytic activity of DNMT3A can be stimulated by DNMT3L or DNMT3B3, two catalytically inactive accessary proteins (20,21). The crystal structure of the methyltransferase domain of DNMT3A and DNMT3L showed a heterotetramer complex of DNMT3L-DNMT3A-DNMT3A-DNMT3L (22). More recently, a cryo-EM structure of DNMT3A-DNMT3B3 heterotetramer bound to a nucleosome revealed a binding between DNMT3B3 catalytic-like domain and the nucleosome acidic patch (23). DNMT3A can also function as homotetramer or oligomer without the accessary DNMT3 proteins (24,25). However, it remains unclear how DNMT3A oligomers interact with the nucleosome core, and whether various DNMT3A-containing higher-order complexes operate at distinct regions of the genome.

In addition to its methyltransferase domain, DNMT3A harbors several regulatory domains that can bind histone PTMs (26). In particular, its PWWP domain recognizes histone H3K36 di- and tri-methylation (H3K36me2/3), which facilitates targeting to gene bodies and promoter-proximal intergenic regions of highly expressed genes (27–31). Missense mutations that disrupt the DNMT3A PWWP-H3K36me2/3 interaction have been reported in patients with Heyn–Sproul– Jackson syndrome or rare neuroendocrine tumors (32–34). Interestingly, cells harboring DNMT3A PWWP mutations showed increased DNMT3A1 localization and DNA hypermethylation at promoter CGIs of Polycomb target genes (32,35–37). Furthermore, the N-terminal region specific to DNMT3A1 long isoform can bind H2AK119Ub *in vitro* (37,38). These studies suggest that imbalanced DNMT3A1 targeting, regulated by the competition between two histone reader activities, may contribute to aberrant DNA hypermethylation of Polycomb target genes.

In this study, we provide structural, biochemical, and cellular analysis to explore the molecular details, functional significance, and cancer relevance of the interaction between DNMT3A1 and H2AK119Ub-nucleosome. We identify critical regions within the DNMT3A1 N-terminus that interact with ubiquitin and the nucleosome acidic patch, which collectively contribute to the genome-wide localization of DNMT3A1 to Polycomb CGIs. We further show that this dual recognition of H2AK119Ub-nucleosome by DNMT3A1 is essential for CGI hypermethylation during tumor progression, and the stable silencing of Polycomb target genes. Together, these results provide insights to the cause, consequence, and therapeutic targeting of cancer-specific CGI hypermethylation.

## Results

### Identifying minimal regions within N-terminus of DNMT3A1 necessary for its localization to H2AK119Ub chromatin

We and others previously reported that disruptions of DNMT3A1 PWWP domain led to its mislocalization to H2AK119Ub-high genomic regions (36–38). Building on these findings, we further explored the nuclear distribution of epitope-tagged DNMT3A1 using immunofluorescence (IF) staining in C3H10T1/2 (10T) cells, a tetraploid female mouse mesenchymal stem cell line. In contrast to the uniform nuclear distribution of wildtype DNMT3A1 (DNMT3A1^WT^), a mutant lacking its PWWP domain (DNMT3A1^ΔPWWP^) localized to discrete foci (Fig. S1A). These foci were also enriched for Xist – a marker for inactive X-chromosome (Xi) - as assessed by RNA-fluorescence *in situ* hybridization (FISH). Consistent with previous studies (39,40), we found that Xi harbors high levels of histone H2AK119Ub readily visible by IF (Fig. S1B). Double deletion of *Ring1a/b*, the catalytic subunits of PRC1, ablated H2AK119Ub and reverted DNMT3A1^ΔPWWP^ distribution to that of DNMT3A1^WT^ (Fig. S1B and S1C). The DNMT3A1^ΔPWWP^ redistribution to Xi was recapitulated by point mutations (W330R, D333N) that compromise the PWWP-H3K36me2 interaction (Fig. S1D). Of note, deleting the PWWP domain of the DNMT3A2 short isoform, which lacks the DNMT3A1 ∼220aa N-terminal region, did not lead to accumulation at Xi (Fig. S1D). Furthermore, swapping the DNMT3A1 N-terminal region into DNMT3B was sufficient to drive DNMT3B^ΔPWWP^ to Xi. These results agree with previous genome-wide studies showing a critical role for the N-terminal region in targeting DNMT3A1 to H2AK119Ub chromatin (37,38). Therefore, we reasoned that IF could serve as a rapid and robust approach to investigate the interaction between DNMT3A1 and H2AK119Ub.

The N-terminus of DNMT3A1 contains a disordered region (aa1-159) and a predicted α-helix (aa160-219) (Fig. S1E). We found that aa160-219 but not aa1-159 of DNMT3A1^ΔPWWP^ were essential for the H2AK119Ub interaction (Fig. S1E), and thus refer to aa160-219 as the ubiquitin-dependent recruitment (UDR) region. To further define the minimal element(s) within the UDR required for the DNMT3A1-H2AK119Ub interaction, we systematically removed four to eight amino acid blocks (denoted ΔR1 to ΔR7) across aa165-212 of DNMT3A1^W330R^, followed by IF for DNMT3A1 (Fig.1 A-C). ΔR1 (aa165-172), ΔR3 (aa181-188) and ΔR4 (aa189-196), but not ΔR2 (aa173-180), ΔR5 (aa197-204) or ΔR7 (aa209-212), fully reversed the Xi accumulation of DNMT3A1^W330R^ (Fig. 1A-B). DNMT3A1^W330R^ ΔR6 (aa205-208) was primarily cytoplasmic, consistent with loss of a nuclear localization signal within this region (Fig. 1A-B) (41). These genetic analyses suggest that multiple regions within the UDR contribute to the DNMT3A1 recognition of H2AK119Ub-modified nucleosome.

**Figure 1.**
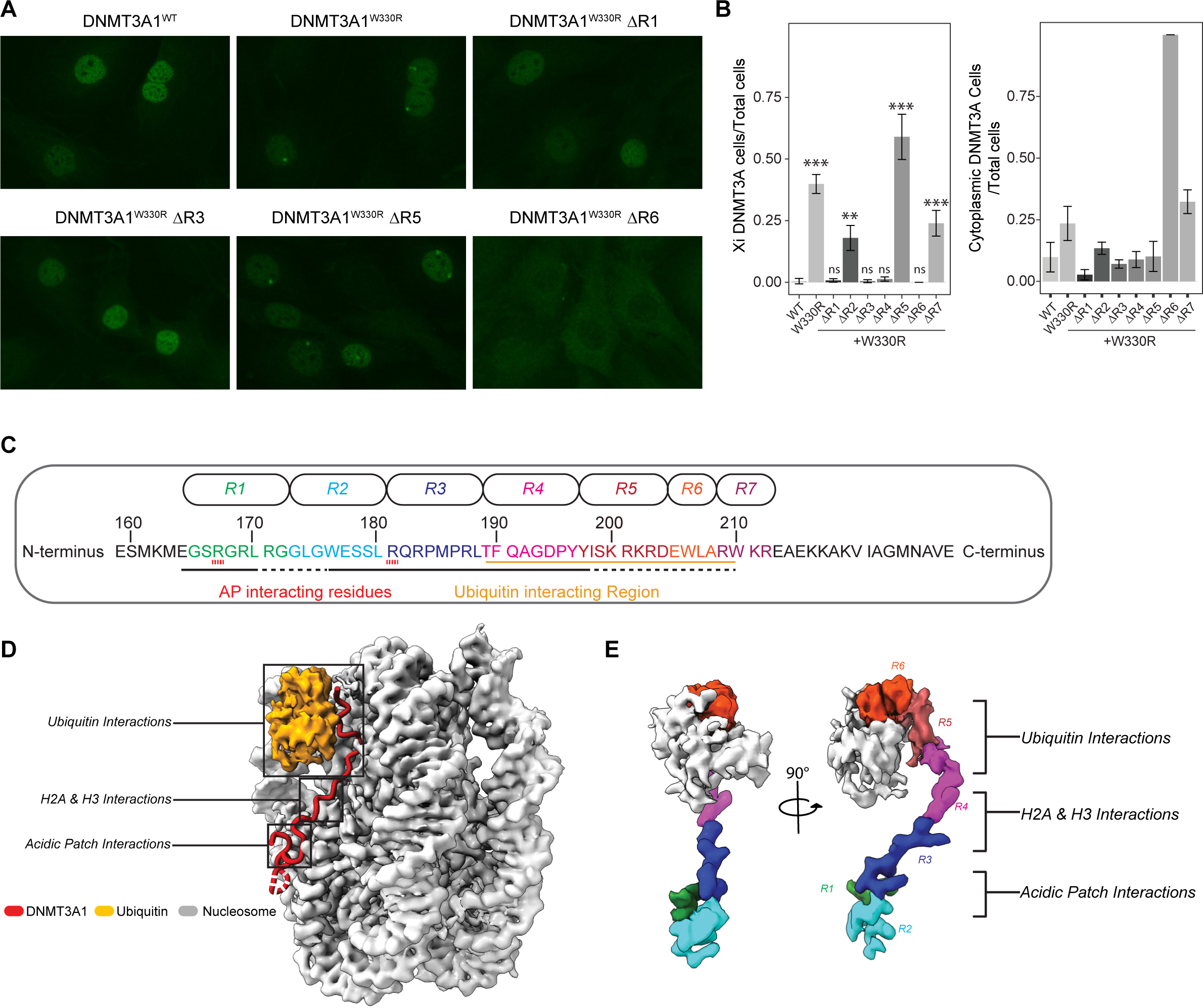
Identification of critical regions within N-terminus of DNMT3A1 for DNMT3A1-H2AK119Ub interaction. **A)** Representative immunofluorescence (IF) staining of FLAG-DNMT3A1 in 10T cells expressing DNMT3A1 wild type, W330R within the PWWP domain, or four-to eight amino acid deletions within the UDR domain (ΔR1, ΔR3, ΔR5 and ΔR6: see Fig.1C for more detail) in the context of DNMT3A1^W330R^. **B)** Quantification of FLAG-DNMT3A1 IF in 10T cells expressing DNMT3A1^WT^, DNMT3A1^W330R^ or DNMT3A1^W330R^ + ΔR1-ΔR7. Left: Ratio of 10T cells displaying DNMT3A1 Xi accumulation compared to all 10T cells. Right: Ratio of 10T cells displaying DNMT3A1 cytoplasmic localization compared to all 10T cells. Error bars represent standard deviation of five replicates. Student t test, ns p>0.05, * p<0.05, ** p<0.01, *** p<0.001. **C)** Amino acid sequence of the DNMT3A1 UDR domain (aa159-228). Regions subject to deletion analysis in Fig. 1A (R1-R7) are indicated at the top. Interaction surfaces for the nucleosome acidic patch and ubiquitin are indicated at the bottom. **D)** Cryo-EM map of DNMT3A1 N-terminal region (aa159-228) bound to H2AK119Ub nucleosome. **E)** Portion of the cryo-EM map highlighting various regions within DNMT3A1 UDR domain (colored as per Fig. 1C) and Ubiquitin, shown at two different views.

### A cryo-EM structure of DNMT3A1 UDR in complex with H2AK119Ub-nucleosome

To gain a detailed understanding of their potential mode(s) of engagement, we determined the structure of the DNMT3A1 UDR (aa159-228) bound to a H2AK119Ub nucleosome at an overall resolution of 2.8 Å (Fig. 1D). This structure shows well-resolved densities for the DNMT3A1 UDR peptide and nucleosome, as well as a density for ubiquitin (Ub) (Fig. S2 and S3). From our cryo-EM map, we can unambiguously assign 33 residues of DNMT3A1 UDR (aa165-197) that span the nucleosome face and make histone contacts (Fig. S4). Density for parts of the peptide interacting with Ub, and the Ub density itself were more mobile, precluding our ability to define specific sidechain interactions (Fig. S4). Furthermore, the Ub interaction region contains several charged residues (DNMT3A1 aa200-203: KRKR) that could also potentially interact with nucleosomal DNA (Fig. 1C-D). Of note, the structure suggested critical regions of interaction between the DNMT3A1 UDR and H2AK119Ub (R6); histones H2A and H3 (R4); and the nucleosome acidic patch (R1 and R3) (Fig. 1E). The structural importance of these blocks is consistent with our deletion analyses (Fig. 1A-C). Our structure also identifies interactions between DNMT3A1 R181 and H2A E61, N90, and E92 of the nucleosome acidic patch (Fig. 2A). While it appears that R181 is the primary acidic patch interacting residue, the UDR peptide folds into a flexible loop, allowing DNMT3A1 R167 to also engage H2A E61 and E64 from the acidic patch. Additionally, DNMT3A1 residues between R167 and R181 engage the nucleosome, further contributing to complex formation.

**Figure 2.**
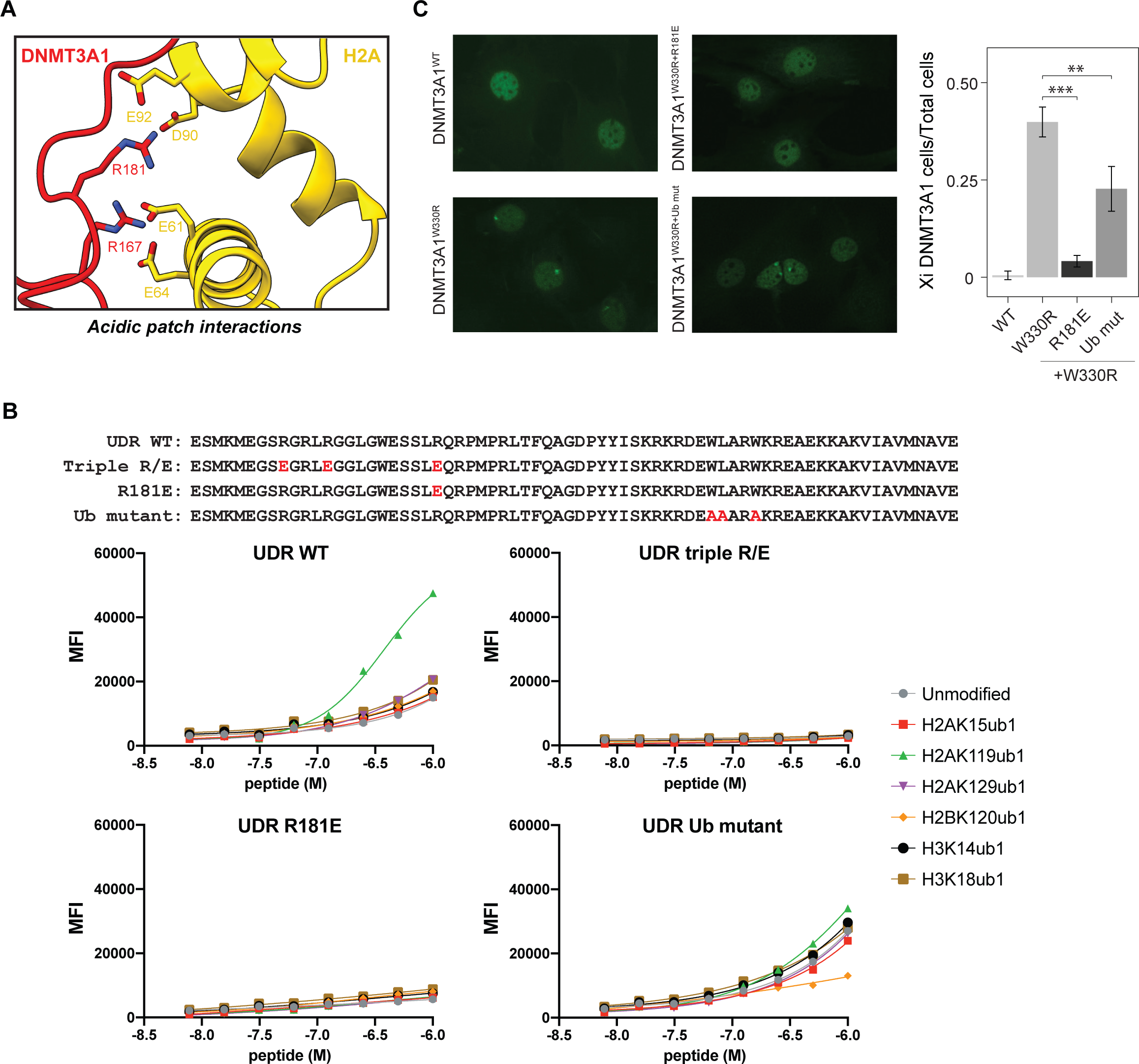
DNMT3A1 UDR domain binds to H2AK119Ub nucleosome via dual recognition of acidic patch and Ubiquitin. **A)** A close-up view of the interactions between the N-terminal region of DNMT3A1 and acidic patch of the nucleosome. **B)** Top: amino acid sequence of the UDR peptide for WT, single acidic patch mutant (R181E), triple acidic patch mutant (R167E, R171E, R181E) and Ub mutant (W207A, L208A and W210A). dCypher-Luminex assay to measure interaction between 6xHis-tagged UDR peptides (WT or mutant as noted: the Queries) and a multiplexed panel of fully defined nucleosomes (the Targets; unmodified, with KUb (H2AK15Ub1, H2AK119Ub1, H2AK129Ub1, H2BK120Ub1, H3K14Ub1, or H2BK18Ub1). **C)** Left: Representative IF of FLAG-DNMT3A1 in 10T cells expressing DNMT3A1^WT^, DNMT3A1^W330R^ (PWWP mutant),DNMT3A1^W330R+R181E^ or DNMT3A1^W330R+Ubmut^. Right: Quantification of FLAG-DNMT3A1 IF in 10T cells expressing DNMT3A1^WT^, DNMT3A1^W330R^, DNMT3A1^W330R+R181E^ or DNMT3A1^W330R+Ub mut^, shown as ratio of 10T cells displaying DNMT3A1 Xi accumulation compared to all 10T cells. Error bars represent standard deviation of five replicates. Student t test, ns p>0.05, * p<0.05, ** p<0.01, *** p<0.001.

### DNMT3A1 UDR-H2AK119Ub nucleosome binding *in vitro* requires both acidic patch and ubiquitin interactions

Based on structural analysis of the DNMT3A1 UDR-H2AK119Ub nucleosome interaction, we generated UDR peptides mutated for residues contacting the acidic patch (R167E/R171E (within R1) and R181E (within R3): aka. UDR^triple R/E^) or ubiquitin (W206A, L207A and W210A within R6: aka. UDR^Ub mut^) (Fig. 1C and 2B). We then used multiplexed dCypher™ Luminex (see Methods) to test interactions between the various forms of DNMT3A1 UDR (the Queries) and a panel of fully defined nucleosomes (the Targets). We first confirmed enhanced interaction between wild-type UDR (UDR^WT^) and H2AK119Ub but no other histone lysine monoubiquitinations tested (H2AK15Ub1, H2AK129Ub1, H2BK120Ub1, H3K14Ub1, H3K18Ub1; Fig. 2B). In contrast, UDR^R181E^ or UDR^triple R/E^ (acid-patch binding mutants) completely lost nucleosome binding (Fig. 2B), while UDR^Ub mut^ largely lost the preference for H2AK119Ub (Fig. 2B). Consistently, IF of 10T cells expressing epitope-tagged DNMT3A1^W330R^ showed that overlaid mutations R181E (*i.e.* DNMT3A1^W330R+R181E^), and to a lesser extent _206_WLARW_210_/AAARA (*i.e.* DNMT3A1^W330R+Ub mut^), reversed Xi accumulation (Fig. 2C). Together, these results demonstrate that interaction between DNMT3A1 and H2AK119Ub requires distinct regions of the UDR to mediate bidentate recognition of the nucleosome acidic patch and ubiquitin.

### Genome-wide localization and methyltransferase activity of DNMT3A1 are regulated by its interface with H2AK119Ub and nucleosome acidic patch

We next determined how the genome-wide targeting and *de novo* methyltransferase activity of DNMT3A1 were impacted by mutations in its UDR or PWWP domains. To this end, we created DNMT quadruple knockout (QKO) mouse embryonic stem cells (mESCs) by targeted deletion of *Dnmt3l* in triple knockout (TKO) mESCs (42) already deficient for *Dnmt3a*, *Dnmt3b* and *Dnmt1* (Fig. S5A). We then re-expressed either wild-type Dnmt3a1 (DNMT3A1^WT^) or alleles carrying mutations in the PWWP domain (DNMT3A1^W330R^), acidic patch interface (DNMT3A1^R181E^), or ubiquitin-binding region (DNMT3A1^Ub mut^) (Fig. S5B). This system enabled us to assess DNMT3A1 enrichment (by ChIP-seq) and *de novo* DNAme (5mC/5hmC by Enzymatic Methyl-seq; EM-seq) in the absence of any additional DNA methyltransferase machinery. As expected, while DNMT3A1^WT^, DNMT3A1^R181E^ and DNMT3A1^Ub mut^ localization and methylation activity positively correlated with H3K36me2, this was not observed with DNMT3A1^W330R^ (Fig. 3A-C; Fig. S5C). Despite the co-localization of DNMT3A1^R181E^ and H3K36me2, we noticed a general decrease in methyltransferase binding and activity compared to DNMT3A1^WT^ (Fig. 3A-C; Fig. S5C). We also analyzed TKO mESCs expressing DNMT3A1 mutants (Fig. S5D), where DNAme was measured by reduced-representation bisulfite sequencing (RRBS). Unlike QKO cells, the overall DNAme level and its positive correlation with H3K36me2 in TKO cells expressing DNMT3A1^R181E^ were similar to that of DNMT3A1^WT^ (compare Fig. 3C and Fig. S5D). This suggested that in the presence of DNMT3L adaptor protein, diminished UDR interaction with the nucleosome acidic patch does not compromise DNMT3A1-mediated methylation at H3K36me2 regions.

**Figure 3.**
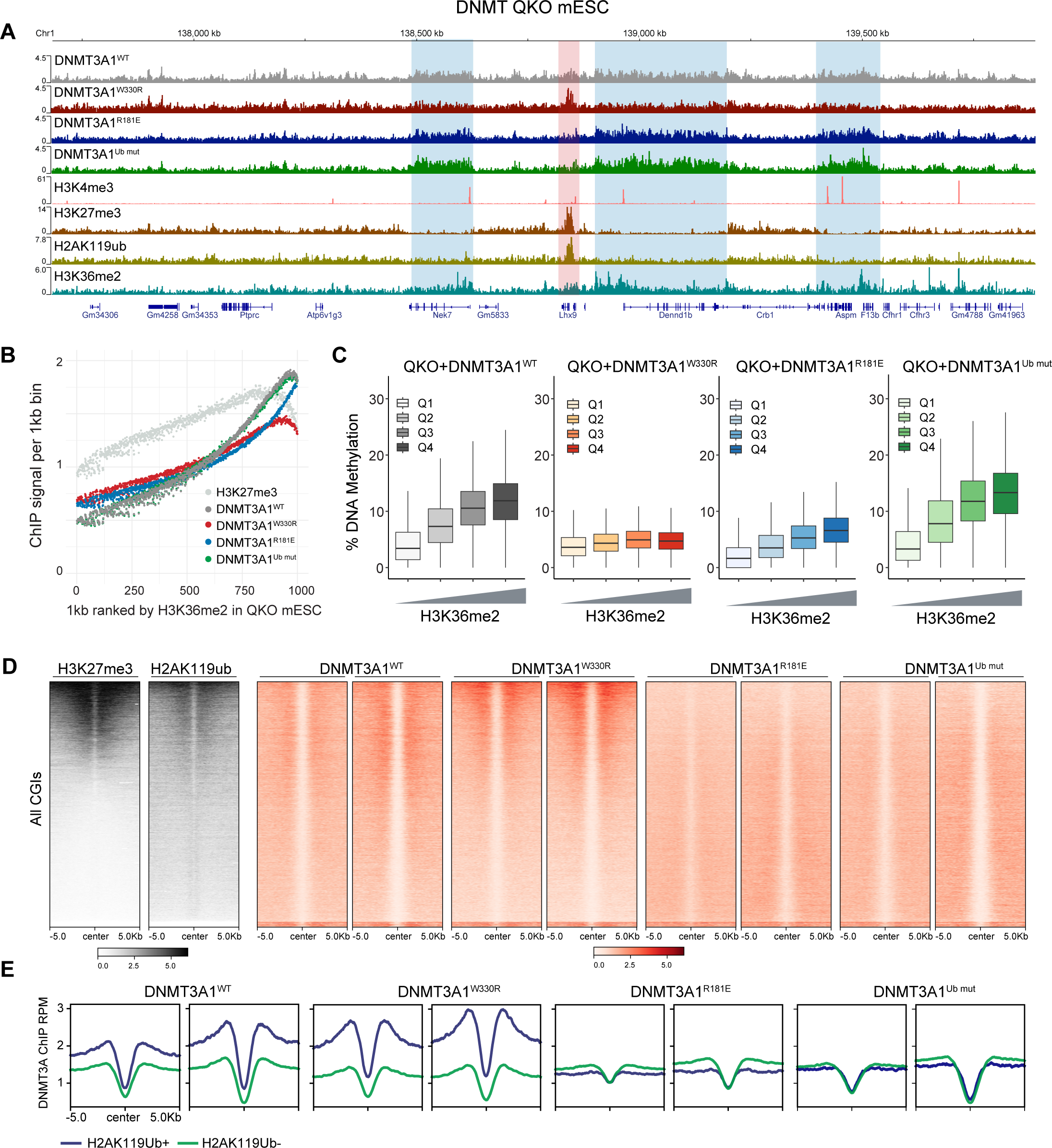
**A)** Genome browser view of DNMT3A1 ChIP–seq reads for DNMT3A1^WT^, DNMT3A1^W330R^, DNMT3A1^R181E^ and DNMT3A1^Ub mut^ QKO mESC at chromosome 1: 137.7–139.8 Mb, GRCm38. Additionally, the Cut and Run reads for the histone modifications H3K4me3, H3K27me3, H2AK119Ub and H3K36me2 in DNMT3A1^WT^ QKO mESC are shown. Genes from the RefSeq database are annotated at the bottom. **B)** Cut and run reads per 1000 averaged 1kb bin for H3K27me3 and ChIP-seq reads per 1000 averaged 1kb bin for DNMT3A1 in DNMT3A1^WT^, DNMT3A1^W330R^, DNMT3A1^R181E^ and DNMT3A1^Ub mut^ QKO mESC. To generate bins, 1kb genomic tiles were ranked by H3K36me2 enrichment in DNMT3A1^WT^ QKO mESC and grouped into 1000 rank-ordered bins. **C)** Percent of DNA methylation (EMseq) in DNMT3A1^WT^, DNMT3A1^W330R^, DNMT3A1^R181E^ and DNMT3A1^Ub mut^ QKO mESC per 10k bins grouped by H3K36me2 CUT&RUN enrichment (Q1 low to Q4 highest). **D)** Enrichment heat map of H3K27me3 and H3K36me2 CUT&RUN reads in DNMT3A1^WT^ QKO mESC and DNMT3A1 ChIP-seq reads in DNMT3A1^WT^, DNMT3A1^W330R^, DNMT3A1^R181E^ and DNMT3A1^Ub mut^ QKO mESC sorted by H3K27me3 centered at CGIs ± 5 kb. **E)** Enrichment plot of ChIP-seq reads of DNMT3A1 in DNMT3A1^WT^, DNMT3A1^W330R^, DNMT3A1^R181E^ and DNMT3A1^Ub mut^ QKO mESC centered at CGIs ± 5 kb and grouped highest 20% H2AK119Ub enriched CGIs (H2AK119Ub+) and lowest 80% H2AK119Ub enriched CGIs (H2AK119Ub-).

We next performed CUT&Run for histone PTMs associated with active (H3K4me3) or Polycomb-regulated (H3K27me3 and H2AK119Ub) CGIs. DNMT3A1^WT^ enrichment was higher in H3K27me3^+^ and H2AK119Ub^+^ CGIs, but negatively correlated with H3K4me3 (Fig. 3D-E, Fig. S5E). This trend was more pronounced for DNMT3A1^W330R^, consistent with the notion that H2AK119Ub competes with H3K36me2 for DNMT3A1 recruitment. In contrast, DNMT3A1^R181E^ and DNMT3A1^Ub mut^ showed no preferential co-localization to H3K27me3^+^ and H2AK119Ub^+^ CGIs (Fig. 3A, D-E and Fig. S5E). These findings suggest that, unlike H3K36me2 regions, interactions between DNMT3A1 UDR and nucleosomal H2AK119Ub / acidic patch are essential for the genome-wide localization of DNMT3A1 to Polycomb-regulated CGIs.

### Redistribution of DNMT3A1 leads to DNA hypermethylation at Polycomb CGIs

Despite wildtype or mutant DNMT3A1 targeting to H3K27me3^+^ and H2AK119Ub^+^ CGIs, we observed minimal DNAme increase at these regions in either QKO or TKO mESCs (Fig. S5F). This is possibly due to high expression of the DNA demethylating TET enzymes in mESCs, which serve to protect CGIs from aberrant DNAme gain (43,44). Supporting this idea, ectopic expression of DNMT3A1^W330R^ in a more differentiated environment, such as 10T mouse mesenchymal stem cells, was sufficient to drive promoter CGI DNA hypermethylation (Fig. 4A-B, Fig. S6A). By integrating RRBS data with CUT&Run analysis of histone PTMs, we found that promoters more likely to become hypermethylated in DNMT3A^W330R^-expressing cells were either H2AK119Ub^+^ or [H2AK119Ub^+^, H3K27me3^+^] in control cells (Fig. S6B, Fig. 4B-C). This DNMT3A1^W330R^-induced hypermethylation of Polycomb CGIs can be reversed by overlaying a UDR mutation that compromises acid patch binding (DNMT3A1^W330R+R181E^), and to a lesser extent one that reduces Ub binding (DNMT3A1^W330R+Ub mut^) (Fig. 4A-B).

**Figure 4.**
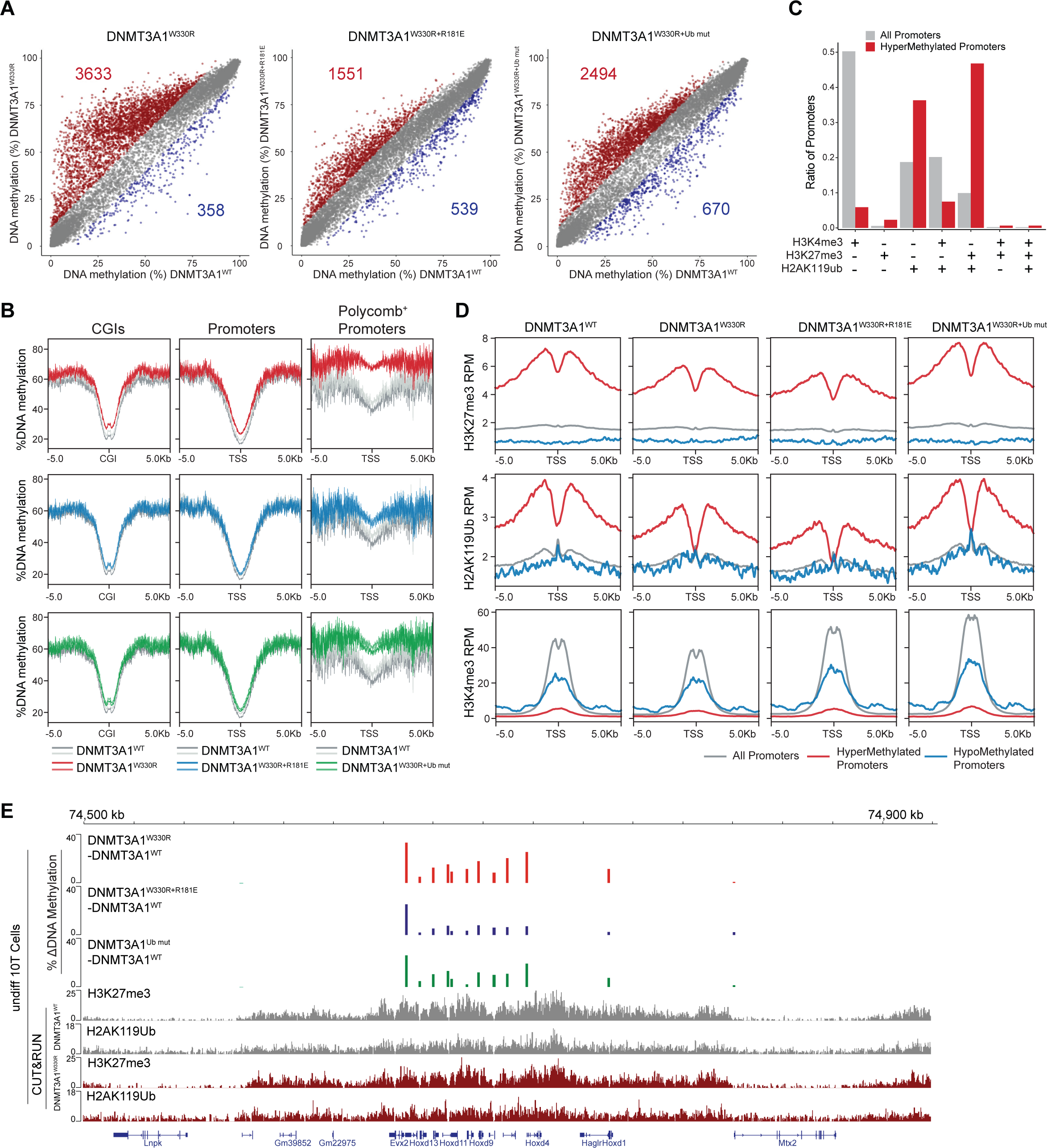
Polycomb promoter hypermethylation following DNMT3A1 redistribution requires DNMT3A1-acidic patch and H2AK119 ubiquitin interaction. **A)** Scatter plots showing % DNAme (measured by RRBS) at all promoters (TSS ± 500bp) in DNMT3A1^W330R^, DNMT3A^W330R+181E^ and DNMT3A1^W330R+Ub mut^ versus DNMT3A1^WT^-expressing 10T cells. Each dot represents a single promoter. Significantly hyper- (>10% and q<0.01) and hypo-methylated (>-10% and q<0.01) promoters are colored in red and blue, respectively. **B)** Metagene plot showing % DNAme over CGIs ± 5 kb (left), promoters (TSS± 5 kb) (middle) and Polycomb+ promoters (H3K27me3^+^ and H2AK119Ub^+^) (right) in DNMT3A1^W330R^, DNMT3A^W330R+R181E^ and DNMT3A1^W330R+Ub mut^ compared to DNMT3A1^WT^ 10T cells. **C)** Barplot showing the representations among all promoters, or 10T DNMT3A^W330R^ hypermethylated promoters, for each group of promoters defined by different histone modificaiton status in control cells as summarized in Fig. S6B. **D)** Metagene plot showing enrichment of H2AK119Ub, H3K27me3 and H3K4me3 CUT&RUN reads from DNMT3A1^WT^, DNMT3A1^W330R^, DNMT3A^W330R+R181E^ and DNMT3A1^W330R+Ub mut^ 10T cells, centered at TSS ± 5 kb and grouped by all, hypermethylated and hypomethylated promoters. **E)** Genome browser view at Chromosome 2, 74.5–75.0Mb, showing differences in % DNA methylation (measured by RRBS) between various DNMT3A1 mutant versus DNMT3A1^WT^ 10T cells. Bottom shows H2AK119Ub, H3K27me3 and H3K4me3 CUT&RUN reads from DNMT3A1^WT^ and DNMT3A1^W330R^ 10T cells. Genes from RefSeq database are annotated at bottom.

We next performed CUT&RUN for H3K4me3, H3K27me3 and H2AK119Ub in various DNMT3A1 backgrounds to further characterize the chromatin state at CGIs that gained DNAme. In DNMT3A1^W330R^ cells, while both genome-wide analysis and inspection of representative regions (*e.g.*, *HoxD* genes) showed modest (∼22%) reductions in H3K27me3 and H2AK119Ub at DNA hypermethylated CGIs, such regions still retained > two-fold higher enrichment of these Polycomb marks relative to all promoters (Fig. 4D-E). Additionally, there was no correlation between the change in H3K27me3 and degree of DNAme increase (Fig. S6C). Therefore, it appears that PWWP mutant-induced DNMT3A1 redistribution leads to aberrant DNAme at Polycomb-regulated CGIs in a UDR-dependent manner. This results in a hybrid chromatin state defined by the co-existence of three repressive epigenetic marks: H3K27me3, H2AK119Ub and DNAme.

### DNA hypermethylation at Polycomb CGIs impairs differentiation-induced transcriptional activation of target genes

We performed RNA-seq to determine if co-existing DNAme and Polycomb PTMs impact gene expression in DNMT3A1 mutant cells. Only a few genes were significantly differentially expressed between 10T cells expressing DNMT3A1^WT^ or DNMT3A1^W330R^ (Fig. 5A), and the upregulated and downregulated genes showed minimal and comparable changes in DNAme (Fig. S6D). Additionally, genes that were hypermethylated in DNMT3A1^W330R^ cells already belonged to the lowest expressed percentiles in DNMT3A1^WT^ cells (Fig. 5B), suggesting they are effectively repressed by Polycomb complexes and the introduction of DNAme has no further silencing effect at the steady state level.

**Figure 5.**
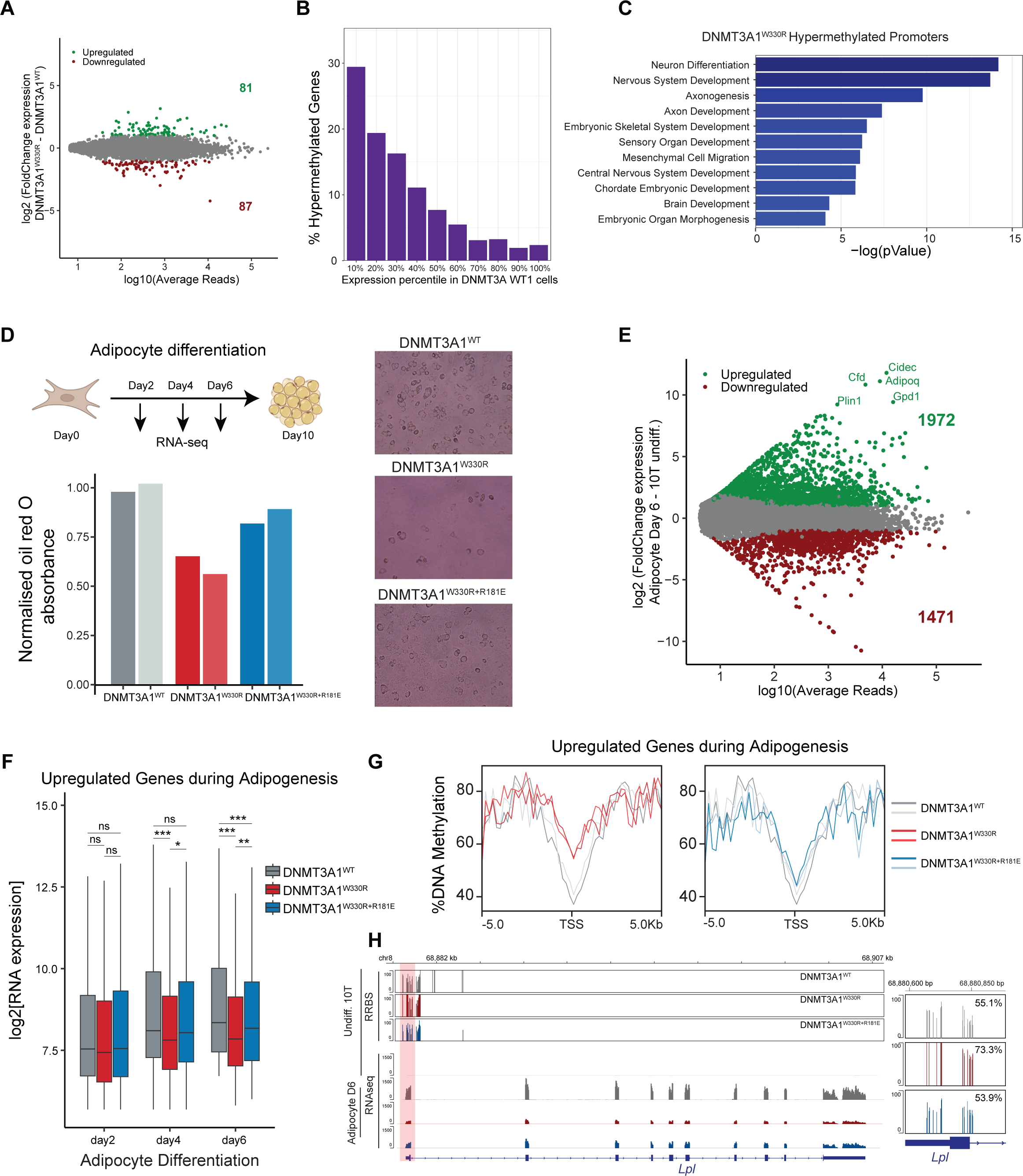
**A)** MA-plot comparing log10 of average normalized reads (RNAseq) per gene in DNMT3A1^W330R^ and DNMT3A1^WT^ 10T (x-axis) vs log2 fold change (LFC) in gene expression (RNAseq) between DNMT3A1^W330R^ and DNMT3A1^WT^ (y-axis). Significantly differently expressed genes are highlighted in green (LFC>2 and q<0.01) or red (LFC<-2 and q<0.01). **B)** Barplot showing distribution of hypermethylated genes (10T DNMT3A1^W330R^ vs 10T DNMT3A1^WT^) into gene expression deciles in DNMT3A1^WT^ 10T. **C)** Barplot of log10 pValues of select GO terms of hypermethylated genes (10T DNMT3A1^W330R^ vs 10T DNMT3A1^WT^). **D)** Top: Schematic of 10T to Adipocyte differentiation over 10 days. Lower: Barplot of normalized Red Oil O absorption in DNMT3A1^WT^, DNMT3A1^W330R^, DNMT3A^W330R+181E^ day 10 Adipocytes. Right: Representative images of Red Oil O staining in DNMT3A1^WT^, DNMT3A1^W330R^, DNMT3A^W330R+181E^ day 10 Adipocytes. **E)** MA-plot comparing log10 of average normalized reads (RNAseq) per gene in DNMT3A1^WT^ day 6 Adipocyte diff. 10T cells and DNMT3A1^WT^ undiff. 10T (x-axis) vs log2 fold change (LFC) in gene expression (RNAseq) between in DNMT3A1^WT^ day 6 Adipocyte diff. 10T cells and DNMT3A1^WT^ undiff. 10T (y-axis). Significantly differently expressed genes are highlighted in green (LFC>2 and FDR<0.01) or red (LFC>-2 and FDR<0.01). **F)** Gene expression (log2 RPM) boxplot of upregulated genes during Adipogenesis (500 genes) in adipocyte differentiated day 2, day 4 and day 6 for DNMT3A1^WT^, DNMT3A1^W330R^, DNMT3A^W330R+181E^ cells. Student t test, ns p>0.05, * p<0.05, ** p<0.01, *** p<0.001. **G)** Enrichment plot showing the distribution of % DNA methylation over the promoters ± 5kb of upregulated genes during adipogenesis for DNMT3A1^WT^, DNMT3A1^W330R^, DNMT3A^W330R+181E^ undiff. 10T cells. **H)** Genome browser view of % DNA methylation (RRBS) of DNMT3A1^WT^, DNMT3A1^W330R^, DNMT3A^W330R+181E^ undiff. 10T cells and gene expression of DNMT3A1^WT^, DNMT3A1^W330R^, DNMT3A^W330R+181E^ day 6 Adipocyte diff. 10T cells over the *Lpl* gene. Right panel: Average DNA methylation shown at the *Lpl* promoter. Genetrack from the RefSeq database.

GO analysis of the hypermethylated genes in DNMT3A1^W330R^ cells revealed enrichment for developmental gene clusters such as “Neuron Differentiation”, “Embryonic Skeletal System Development”, and “Embryonic Organ Morphogenesis” (Fig. 5C). Therefore, we examined the dynamics of gene expression during cell differentiation. We induced adipocyte differentiation of DNMT3A1 wildtype or mutant 10T cells as previously (45) for 10 days, with total RNA collected for RNA-seq on days two, four and six of differentiation (Fig. 5D). Red Oil O staining of day 10 adipocytes found visibly reduced differentiation of DNMT3A1^W330R^ cells (Fig. 5D, 61% of DNMT3A1^WT^), and a partial rescue of differentiation in double mutant DNMT3A1^W330R+R181E^ cells (85% of DNMT3A1^WT^). In DNMT3A1^WT^ cells, 3443 genes were differentially expressed comparing day six vs. day 0 of adipocyte differentiation (1972 upregulated / 1471 downregulated) (Fig. 5E). As expected, GO analysis of the upregulated genes day six post-differentiation showed enrichment for adipocyte specific groups (*e.g.*, “Fatty Acid Catabolic Process”, “Fatty Acid Metabolic Process”) (Fig. S6E). Consistent with Red Oil O staining, many adipogenesis-associated genes (46,47), such as *Cfd*, *Cidec*, and *Adipoq*, failed to be activated by day six in DNMT3A1^W330R^ but not DNMT3A1^W330R+R181E^ cells (Fig. S6F). Focusing on the top 500 genes upregulated during adipogenesis, we profiled their expression kinetics through differentiation (days two - six) and observed a significantly delayed increase in DNMT3A^W330R^ cells with a significant rescue observed for DNMT3A^W330R+R181E^ cells (Fig. 5F).

Next, we examined the promoter methylation state of genes upregulated during adipogenesis, using the RRBS data of undifferentiated 10T cells. Compared to all gene promoters, those upregulated during adipogenesis gained a higher degree of DNAme in DNMT3A1^W330R^ cells, with this partially reversed in DNMT3A^W330R+R181E^ cells (Fig. 5G & Fig. S6G). This is especially evident for adipogenesis regulators known to be sensitive to DNAme regulation, such as *Slc27a1* (*Fatp1*), *Klf15* and *Lpl* (Fig. 5H & Fig. S6H) (46). Therefore, the incorporation of DNAme into promoter CGIs harboring Polycomb marks did not affect basal silencing, but limited their potential to become activated upon induction. In the context of adipocyte differentiation, impaired expression of hypermethylated master regulators could have a downstream effect on the less DNAme sensitive (*e.g.*, low promoter CG density) adipogenesis factors (such as *Cfd*, *Cidec*, and *Adipoq*), causing a systematic delay to adipocyte commitment.

### Cancer-associated Polycomb CGI hypermethylation requires DNMT3A1-acidic patch interaction

We next sought to explore the cancer relevance of our findings. Analysis of patient tumor samples suggests that CGIs that gain aberrant DNAme are more likely to be enriched for H3K27me3 in human embryonic or adult stem cells (11–14). However, this correlation has not been mechanistically dissected in a controlled experimental setting, and the potential involvement of DNMT3A not explored. To this end, we employed a well-established carcinogen-induced esophagus squamous cell carcinoma (ESCC) mouse model (48). We generated organoid cultures from either esophagus harvested from normal mice (EN), or ESCC developed in mice after 16 weeks treatment with the carcinogen 4-Nitroquinoline 1-oxide (4NQO) (Fig. 6A). These organoid cultures have been extensively characterized and faithfully recapitulate the genomic and histological features of human normal esophagus and ESCC (48,49). DNAme profiling by RRBS, as well as CUT&Tag for H3K4me3, H3K27me3 and H2AK119Ub, were performed in six independently generated organoid lines (3 x EN; 3 x ESCC). We also performed RRBS in EN organoids after introducing DNMT3A1^WT^, DNMT3A1^W330R^ and DNMT3A1^W330R+R181E^ (Fig. 6A). Using these datasets, we examined: (i) if H2AK119Ub/H3K27me3 enrichment in esophageal progenitors indeed predicts CGI DNA hypermethylation in tumors arising from these cells-of-origin; and, (ii) if redistribution of DNMT3A1 to Polycomb CGIs through impaired PWWP reader function, in a UDR-dependent manner, is sufficient to recapitulate the cancer-specific DNA hypermethylation signature.

**Figure 6.**
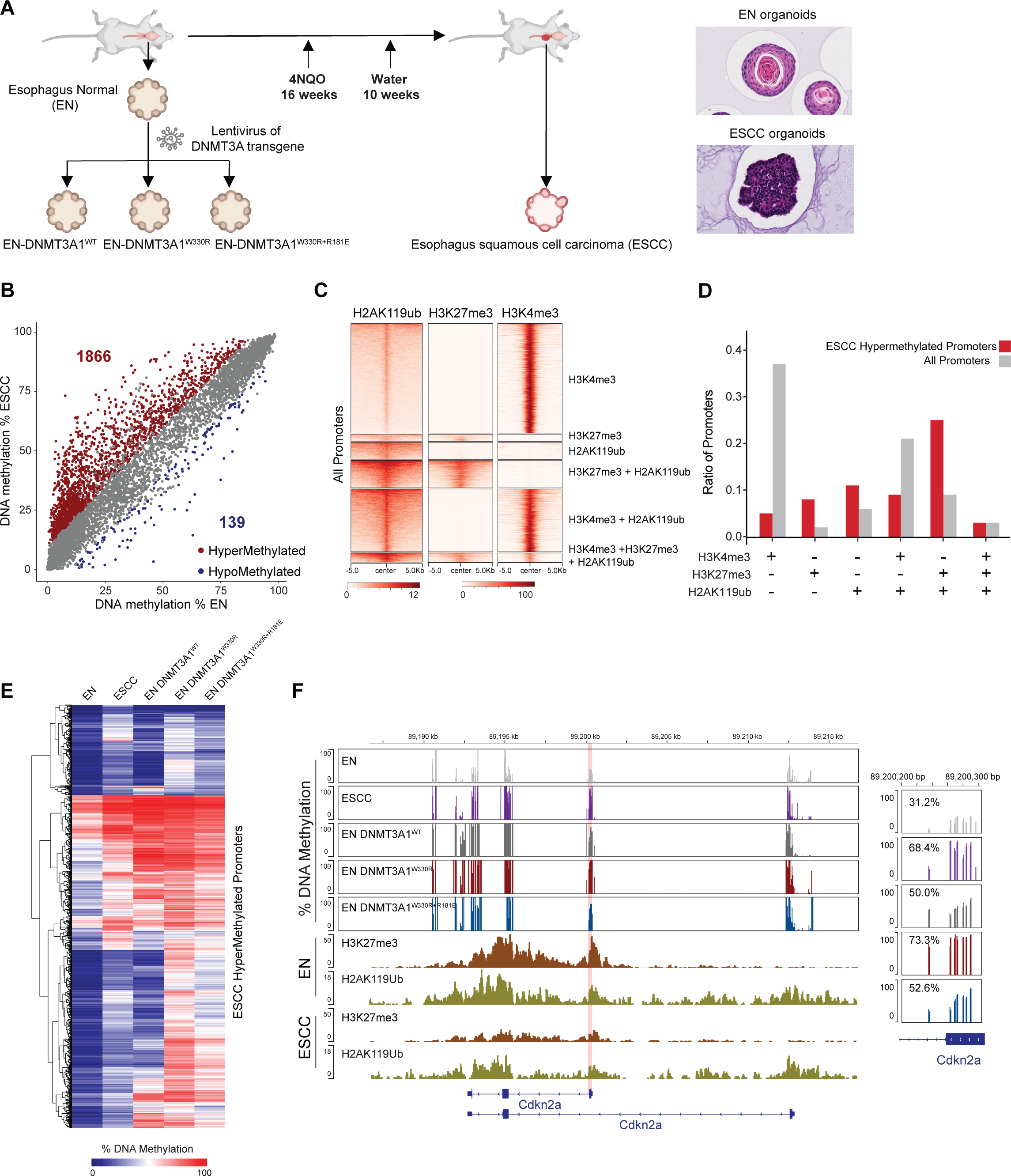
Cancer-associated Polycomb CGI hypermethylation requires DNMT3A1-acidic patch interaction. **A)** Schematic of normal esophagus (EN) and 4NQO-induced esophageal squamous cell carcinoma (ESCC) organoid collection. Right: Representative images of EN and ESCC organoid histology. **B)** Scatter plots showing % DNAme (measured by RRBS) at all promoters (TSS ± 500bp) in ESCC versus EN organoids. Each dot represents a single promoter. Significantly hyper- (>10% and q<0.01) and hypo-methylated (>-10% and q<0.01) promoters are colored in red and blue, respectively. **C)** Heat map showing enrichment of H3K27me3, H2AK119Ub and H3K4me3 CUT&TAG reads at all promoters (TSS± 5kb) in EN organoids. Promoters are classified into six groups: H3K4me3^+^; H3K27me3^+^; H2AK119Ub^+^; H3K4me3^+^ and H2AK119Ub^+^; H3K27me3^+^ and H2AK119Ub^+^; H3K4me3^+^ and H3K27me3^+^ and H2AK119Ub^+^. **D)** Barplots showing the representations among all promoters, or ESCC hypermethylated promoters, for each of the six groups of promoters as defined in Fig. 6C. **E)** Heatmap showing % promoter methylation (measured by RRBS) in EN organoids, ESCC organoids, and EN organoids expressing DNMT3A1^WT^, DNMT3A1^W330R^, DNMT3A1^W330R+R181E^ for all ESCC hypermethylated promoters. **F)** Genome browser view of *Cdkn2a* gene (chromosome 4: 89.187 Mb -89.216 Mb, GRCm39), showing %DNAme (measured by RRBS) in EN organoids, ESCC organoids, and EN organoids expressing DNMT3A1^WT^, DNMT3A1^W330R^, DNMT3A1^W330R+R181E^. Bottom shows H3K27me3 and H2AK119Ub CUT&TAG reads in EN and ESCC organoids. Right panel: Average DNA methylation shown at the *Cdkn2a* promoter (chromosome 4: 89.200.200 bp -89.200.300 bp, GRCm39). Genes from RefSeq database are annotated at the bottom.

RRBS revealed widespread hypermethylation of >1800 promoters and limited hypomethylation of ∼150 promoters in ESCC compared to EN organoids (Fig. 6B), despite comparable global DNAme levels (Fig. S7A). We classified the promoters based on their levels of H2AK119Ub, H3K27me3 and H3K4me3 in EN organoids (Fig. 6C), and found that those with either H2AK119Ub and H3K27me3, or H2AK119Ub alone, were more prone to gaining DNAme in ESCC organoids (Fig. 6D). In contrast, H3K4me3^+^ promoters were more resistant to cancer-associated DNA hypermethylation (Fig. 6D). These results provide strong support for the notion that Polycomb activity in normal tissue predisposes promoter CGIs to hypermethylation during oncogenesis. Similar to 10T cells, DNA hypermethylated promoters in ESCC organoids showed reduced but higher-than-genome average levels of H3K27me3, possibly signifying H3K27me3 and DNAme co-occupancy at these regions (Fig. S7B). However, we observed that compared to 10T cells, the ESCC-associated decreases in H3K27me3 at Polycomb CGIs were stronger and better correlated with the DNAme increase (Fig. S7B).

Of note, expressing DNMT3A1^W330R^ in EN organoids largely recapitulated the hypermethylation signature of ESCC organoids (Fig. 6E, Fig. S7C-D). Over half of the ESCC hypermethylated promoters were also hypermethylated in DNMT3A1^W330R^ EN organoids, including classic tumor suppressor genes such as *Cdkn2a* (50) (Fig. 6F). Furthermore, using DNAme data from TCGA (51), we identified sets of hypermethylated promoters in human esophageal (ESCA) and head and neck (HNSC) cancers. Over 75% of the ESCA and HNSC hypermethylated promoters were also hypermethylated in the DNMT3A1^W330R^ EN organoids (Fig. S7E). Importantly, the DNMT3A1^W330R^ induced aberrant hypermethylation phenotype was markedly reversed in DNMT3A1^W330R+R181E^ organoids (Fig. 6E-F; Fig. S7C-D), suggesting that interactions between DNMT3A1 UDR and nucleosomal H2AK119Ub / acidic patch play a critical role in establishing cancer-associated CGI hypermethylation.

## Discussion

In this study, we provide structural, biochemical and genetic evidence that DNMT3A1 localization to Polycomb CGIs requires binding by its N-terminal UDR to both nucleosomal H2AK119Ub and the acidic patch. This targeting mechanism competes with the DNMT3A1 PWWP-H3K36me2/3 interaction to regulate genome-wide distribution of the methyltransferase. Furthermore, perturbation of the normal competition can lead to the CGI hypermethylation phenotype commonly found in cancer cells. These findings advance our mechanistic understanding of DNMT3A1 recruitment and regulation, how mis-targeting may contribute to tumor development, and represent a therapeutic opportunity.

DNMT3A1 does not have a classical ubiquitin-binding domain, and is not reported to bind free ubiquitin, but still shows preference for H2AK119Ub. This is not the first instance where H2AK119Ub is implicated in pathological mis-localization of chromatin regulators in cancer. The fusion oncoprotein SS18-SSX1 delocalizes BAF chromatin-remodeling complex from promoters/enhancers towards Polycomb-repressed regions marked by H2AK119Ub (52). This targeting is facilitated by SSX1, which binds H2AK119Ub nucleosomes despite lacking a canonical ubiquitin-binding domain and having no affinity for free ubiquitin. Selective binding is accomplished by SSX1 docking to the nucleosome acidic patch and establishing interactions with ubiquitin, much like in the case of DNMT3A1. However, the interactions with ubiquitin differ between these two structures. In SS18-SSX1 interactions extend from the acidic patch through H2A α2 and α3 helices to a surface created by histone H3 and H2AK119Ub (53). In DNMT3A1 the interactions instead extend from the acidic patch through α2, a short loop between α3 and αC οf H2A, and ends between α2 οf H3 and ubiquitin. Despite these distinctions, the general mode of interaction is preserved. These data raise important questions: why are H2AK119Ub nucleosomes uniquely capable in their aberrant delocalization of chromatin complexes (DNMT3A1 PWWP mutant, SS18-SSX1 fusion), and if so, are other complexes mistargeted to Polycomb loci using similar mechanisms?

Short isoform DNMT3A2 lacking the DNMT3A1 N-terminus, including the UDR region, is predominantly expressed in early embryonic development (54). This coincides with the expression of non-catalytic DNMT3L (55), and the resulting DNMT3A2-DNMT3L heterotetramer establishes genome-wide DNAme (22,56,57). In this context, the catalytic-like domains of accessory DNMT3 proteins such as DNMT3B3 or DNMT3L can stabilize DNMT3A2 and/or provide interaction with the nucleosome acidic patch (23) in the absence of UDR. DNMT3A1 becomes the dominant DNMT3A isoform around the same development window when DNMT3L expression decreases (58). We speculate this switch allows DNMT3A1 targeting to Polycomb-regulated genes for *de novo* DNA methylation and long-term silencing (59). For example, a group of maternal genes are non-canonically imprinted by Polycomb activity before they acquire DNAme in extraembryonic tissues after implantation (60). The ability of the DNMT3A1 UDR region to interact with the nucleosome acidic patch may allow the enzyme to act “solo” as a homotetramer or another type of oligomer in these regions without DNMT3L or DNMT3B3. Of note, deletion of the mouse DNMT3A1 N-terminus results in abnormal post-natal development (38). It will be interesting to examine if the growth delay and behavior deficits in these mice are associated with a dysregulated switch from Polycomb to DNAme at certain developmental genes.

At the cellular level, perturbing interaction between the DNMT3A1 UDR and the nucleosome acidic patch (R181E) reduced enzyme binding / activity towards H3K36me2 domains, but essentially abolished localization to H2AK119Ub-marked CGIs. Two non-mutually exclusive mechanisms could contribute to this differential impact. First, the UDR-acidic patch interaction may play a more critical role in stabilizing the DNMT3A1 engagement with H2AK119Ub over H3K36me2 nucleosomes. This could be due to the dual affinity of the PWWP domain for H3K36me2 and DNA, which may overcome the need for acidic patch binding (27). Second, as discussed above, of all DNMT proteins and isoforms only DNMT3A1 has a UDR to mediate H2AK119Ub binding, and is thus most effective *in vivo* as a homo-tetramer or homo-oligomer to engage Polycomb-marked promoters (43,61). In contrast, DNMT3L and DNMT3B3 harbor PWWP domains which enable their targeting to H3K36me2 regions in complex with DNMT3A1, thereby providing additional contacts with the nucleosome acidic patch. This is observed in ESCs where DNMT3L co-localizes with and directs DNMT3A towards H3K36me2/3-marked gene bodies (62). Indeed, we showed that in TKO mESCs (*i.e.* presence of DNMT3L), R181E mutation did not reduce DNMT3A1’s activity at H3K36me2 domains (Figure S5D, compared to QKO data in Fig. 3C). Further structure-function analysis is warranted to test these hypotheses.

We and others have previously shown that mutations in TET DNA demethylases, or inhibition by the “oncometabolite” 2-hydroxyglutatrate produced by gain-of-function mutant isocitrate dehydrogenase (IDH), can establish CGI hypermethylation (63–67). These findings are consistent with the notion that TET enzymes act as guardians to protect CGIs from aberrant DNAme. However, how *de novo* DNA methylating activity is targeted to CGIs, particularly those enriched for Polycomb marks, remains elusive (14). An early model proposes that DNMT3 mislocalization in cancer may be facilitated by sequence-specific transcription factors (68,69). However, the ectopic insertion of CGIs in cancer cells does not lead to their hypermethylation, arguing against a *cis*-element mechanism (70). Using organoids isolated from a well-established ESCC model, we could rigorously test and confirm the association between H2AK119Ub/H3K27me3 enrichment in tissue progenitor cells, and CGI hypermethylation in tumors arising from these cells-of-origin. In addition, our results using normal organoids expressing various DNMT3A1 mutants suggest that perturbed reader function of the PWWP domain facilitates cancer-associated CGI hypermethylation in a UDR-dependent manner. These findings provide a unifying model for the establishment of Polycomb CGI DNA hypermethylation in cancer. In normal somatic tissues, a subpopulation of DNMT3A1 can localize to Polycomb CGI via the UDR-H2AK119Ub interaction, but its methyltransferase activity is countered by TET enzymes. In cancer cells, DNA hypermethylation at these regions can result from either insufficient TET activity, or increased binding of DNMT3A1 redistributed from H3K36me2/3 domains. Notably, missense mutations that impair the PWWP-H3K36me2 interaction are only found in rare types of neuroendocrine tumors (31,34), suggesting that non-genetic mechanisms, such as PTMs within the DNMT3A1 PWWP domain, may contribute more generally to cancer-specific CGI hypermethylation.

Catalytic inhibitors of DNMT (*e.g.,* decitabine) have been approved to treat myeloid malignancies. However, their use in solid tumors is limited in part by toxic side effects, as these inhibitors induce genome-wide demethylation that is also deleterious to normal cells. Our structure-function studies pave the way for developing inhibitors that target the bidentate interaction between DNMT3A1 and the H2AK119Ub nucleosome, which would be expected to specifically prevent or reverse CGI hypermethylation in cancer cells, while preserving the DNA methylome in normal tissues.

## METHODS

### Expressing DNMT3A1 mutants in 10T cells

cDNA of mouse Dnmt3A1^WT^ and DnmtA1^W326R^ (corresponding to human W330R) previously cloned into pCDH-EF1-MCS-Neo (System Biosciences) with an N-terminal FLAG-HA epitope tag(37) were used with standard mutagenesis to generate deletions within UDR (ΔR1, aa161-168; ΔR2, aa169-176; ΔR3, aa177-184; ΔR4, aa185-192; ΔR5, aa193-200; ΔR6, aa201-204; ΔR7, aa205-208: corresponding to human ΔR1 (aa165-172), ΔR2 (aa173-180), ΔR3 (aa181-188), ΔR4 (aa189-196), ΔR5 (aa197-204), ΔR6 (aa205-208), ΔR7 (aa209-212)), the acidic patch mutation R177E (corresponding to human R181E), and the Ub mutant (W202A, L203A and W206A; corresponding to human W206A, L207A and W210A). Primers (Table 1) were designed using NEBbaseChanger tool and desired mutations confirmed by Sanger sequencing. To generate concentrated lentivirus, DNMT3A1 mutant vectors were co-transfected with helper plasmids psPAX2 and pVSVG into HEK293T. After 72h supernatant was collected and concentrated using the PEG-it system as per manufacturer’s instructions (Systems Biosciences).

C3H10T1/2 mouse fibroblast cells (10T; ATCC), were cultured in Dulbecco’s modified Eagle’s medium (DMEM, Invitrogen) with 10% fetal bovine serum (FBS, Gemini). To generate transgenic cell lines expressing DNMT3A1 mutants, 10T cells were transduced with concentrated lentivirus. 48 h after transduction cells were selected under G418 (1 mg/ml) for one week. Methylation analysis was done after 21 days in culture.

### CRISPR-KO to generate QKO mESC

Single guide (sgRNAs) directed against mouse Dnmt3L were cloned into px458. Dnmt3L KO clones (QKO) were generated by transfecting TKO J1 mESC (42) with sgRNA-containing px458 using Lipofectamine 3000 (Invitrogen) and sorting GFP+ cells after 72h. For single clone isolation, transfected cells were seeded at low density (1,000 cells per 9.6 cm^2^) for expansion, and successful knockout lines confirmed by immunoblot (to DNMT3L) and sequencing over the sgRNA target location (analysis with Tracking of Indels by Decomposition tool; TIDE) (71).

### Expressing DNMT3A1 mutants in TKO and QKO mESC

TKO and QKO mESC were cultured in Serum/LIF media (DMEM-KO, 15% FBS, 1000 U/ml murine recombinant LIF (Millipore Sigma #ESG1106), 1× nonessential amino acids (100x, Gibco #11140), 1× GlutaMax (100x Gibco #35050-061) and 0.1 mM 2-mercaptoethanol (Gibco)).

The pCDH-EF1-MCS-Neo vectors carrying Dnmt3A1^WT^, Dnmt3A1^W330R^, Dnmt3A1^R181E^ or Dnmt3A1^Ub mut^ were digested with XbaI and BamHI and cloned to piggyBac backbone vector with GFP visualization marker (pPB-Dnmt3A1-GFP). These were co-transfected with piggybac transpose vector using Lipofectamine 3000 (Invitrogen) to the QKO mESC cell line. GFP+ cells were sorted after 7 days and stable DNMT3A QKO mESC lines were confirmed by flow analysis and immunoblotting for DNMT3A, and sequence confirmation of the appropriate allele. Methylation analysis and DNMT3A ChIP were done after 21 days in culture.

### Adipocyte differentiation

10T cells were seeded at 500,000 cells per 9.6cm^2^ (in DMEM + 10% FBS) and the next day changed to adipocyte differentiation media (DMEM, 10% FBS, 0.5 mM isobutylmethylxanthine, 1 μM dexamethasone, 5 μg/mL insulin and 5 μM troglitazone). After two days media was changed to DMEM +10% FBS + 5 μg/mL insulin and changed every two days until day 10 of adipocyte differentiation. To quantify adipocyte differentiation with Oil Red-O staining, cells were directly cross-linked (3.7% paraformaldehyde for two min) on a plate after media removal and PBS wash. After a water wash, Oil Red-O solution (1ml per 9.6cm^2^; Sigma, #01391) was added and plates incubated at room temperature (RT°) for one hour. Oil Red-O solution was removed, the plate dried and 1ml 100% isopropanol added for 20 minutes at RT°. 200 μl of this isopropanol was used to measure absorbance at 520nm in triplicates on a 96 well plate reader.

### ChIP-seq

Cross-linking ChIP in QKO was done using 2 × 10^7^ cells per immunoprecipitation. Cells were directly cross-linked (1% paraformaldehyde for 10 min at RT° with gentle shaking) on a plate after media removal and PBS wash, and reactions quenched with glycine (added to 125 mM for five min at RT). Cells were washed with cold PBS containing protease inhibitors (Roche, #5056489001), scraped off the plates, pelleted by centrifugation (1200 rpm, 4°C), and flash frozen.

To prepare [antibody: bead] complexes 75ul of Protein A Dynabeads (Invitrogen, #10002D) per ChIP sample were washed three times with PBS + 0.01% Tween20 (collecting on a magnet at each step) and 10ul of anti-DNMT3A (Cell Signaling, #3598) added. Binding reactions were incubated overnight (4 °C under rotation), washed three times with PBS + 0.01% Tween20 (collecting on a magnet at each step), and resuspended in 100ul of LB3.

To obtain a soluble chromatin extract cell pellets were thawed, resuspended in 1 ml LB1 (50 mM HEPES, 140 mM NaCl, 1 mM EDTA, 10% glycerol, 0.5% NP-40, 0.25% Triton X-100 and 1× protease inhibitor (Roche, #5056489001)) and placed on a rotator (10 min at 4 °C). Samples were collected by centrifugation (1350 g for 5 min), resuspended in 1 ml LB2 (10 mM Tris-HCl pH 8.0, 200 mM NaCl, 1 mM EDTA, 0.5 mM EGTA and protease inhibitor), and placed on a rotator (10 min at 4 °C). Samples were collected by centrifugation (1350 g for 5 min), resuspended in 1 ml LB3 (10 mM Tris-HCl pH 8.0, 100 mM NaCl, 1 mM EDTA, 0.5 mM EGTA, 0.1% Na-deoxycholate, 0.5% N-lauroylsarcosine, 1% Triton X-100 and protease inhibitor). To prepare chromatin, extracts were sonicated for 20 min using a Covaris M220 focused ultrasonicator (peak power 140, duty factor 5, cycles/burst 200) and centrifuged at 4°C for 20min to remove cell debris.

Aliquots of sonicated chromatin (5% each) were removed as Input DNA or to confirm effective sonication (by resolving on an agarose gel and staining for DNA). The remaining material was incubated (overnight at 4 °C on a rotator) with 100ul of anti-DNMT3A bound magnetic beads. Beads were then sequentially collected on a magnet / washed with low-salt buffer (150 mM NaCl; 0.1% SDS; 1% Triton X-100; 1 mM EDTA, 50 mM Tris-HCl), high salt buffer (500 mM NaCl; 0.1% SDS; 1% Triton X-100; 1 mM EDTA, 50 mM Tris-HCl), LiCl buffer (150 mM LiCl; 0.5% Na deoxycholate; 0.1% SDS; 1% Nonidet P-40; 1 mM EDTA, 50 mM Tris-HCl) and TEN buffer (1 mM EDTA, 10 mM Tris-HCl, 50mM NaCl). After the final wash and 3 min centrifugation beads were resuspended in elution buffer (1% SDS, 50 mM Tris-HCl pH 8.0, 10 mM EDTA and 200 mM NaCl) and incubated for 30 min at 65 °C. After brief centrifugation (20,000xg) the supernatant was collected, and cross-links reversed by overnight incubation at 65 °C. The eluate was then sequentially treated with RNase A (Thermo Scientific; 1 h at 37 °C) and Proteinase K (Roche; 1 h at 55 °C) and DNA recovered using PCR purification kit (Qiagen).

For ChIP–seq, libraries were prepared with NEBNext Ultra II Library Prep Kit reagents as per manufacturer’s instructions. Libraries were sequenced on an Illumina NextSeq 550 to generate 75-bp single-end reads.

### CUT&Tag (72)

2.5 × 10^5^ cells were collected and washed with 1 ml of wash buffer (20 mM HEPES pH 7.5, 150 mM NaCl, 0.5 mM Spermidine (ThermoFisher, #13274)) before being resuspended in 1ml of wash buffer. Concanavalin A-coated magnetic beads (Bangs Laboratories, #BP531) were washed twice with 1 ml binding buffer (20 mM HEPES pH 7.5, 10 mM KCl, 1 mM MnCl_2_, 1 mM CaCl_2_). 10 μl of washed magnetic beads were added to each sample of 2.5 × 10^5^ cells and incubated with rotation at RT° for 15 min. Bead-bound cells were collected on a magnetic rack and resuspended in 100 μl of antibody buffer (Wash buffer + 0.05% digitonin (Sigma-Aldrich, #D141), 2 mM EDTA) and incubated with a primary antibody at 4 °C overnight on nutator. After washing (with Wash buffer + 0.05% Digitonin), bead-bound cells were incubated with 1 μl anti-rabbit IgG secondary (Antibodies-online, #ABIN101961) and 2 μl pA-Tn5 adapter complex in 100 μl Dig-300 buffer (Wash buffer + 300 mM NaCl, 0.01% Digitonin) at RT for 1 h on nutator. Cells were washed three times with 1ml Dig-300 buffer, resuspended in 250 μl Tagmentation buffer (Dig-300 buffer + 10 mM MgCl_2_), and incubated at 37 °C for 1 h. To each sample, 10 μl of 0.5 M EDTA, 3 μl of 10% SDS and 5 μl of 10 mg/ml Proteinase K (Roche, #31158) were added and incubated at 50 °C for 1 h. DNA was purified using PCR purification kit (Qiagen) and eluted with 25 μl ddH_2_O.

For library amplification, 2 μL i5 unique index primer (10 μM), 2 μL i7 unique index primer (10 μM) and 25 μL NEBNext High-Fidelity 2X PCR Master Mix (NEB) were added to 21 μl of purified CUT&TAG DNA, and the mix subjected to PCR (72 °C, 5 min; 98 °C, 30 sec; 12 cycles of 98 °C, 10 sec and 63 °C, 10 sec; 72 °C, 1 min and hold at 10 °C). To purify PCR products, 1.1× volumes (55 μl) of Ampure XP beads (Beckman Coulter, #A63880) were added and incubated for 10 min at RT. Libraries were washed twice with 80% ethanol and eluted in 20 μl of 10 mM Tris-HCl, pH 8. Libraries were sequenced on an Illumina NextSeq 550 to generate 75-bp paired-end reads.

### CUT&RUN

CUTANA^TM^ CUT&RUN was performed with 10T and QKO samples on an automated protocol (autoCUT&RUN) derived from those previously described (73–75). In brief, for each CUT&RUN reaction 500K cells [from 5 million cells/mL prepared in FBS with 10% DMSO] were dispensed to individual wells of a 96-well plate, immobilized onto Concanavalin-A beads (Con-A; EpiCypher #21-1401), and incubated overnight (4°C) with 0.5 µg of antibody (IgG, H3K4me3, H3K27me3, H3K36me2, H2AK119Ub1) (all antibodies validated to histone post-translation modification (PTM)-defined SNAP-ChIP nucleosome standards as previously (76)). pAG-MNase (EpiCypher #15-1016) was added / activated (2 hours @ 4°C) and CUT&RUN enriched DNA purified using Serapure beads after mixing at 2:1 (Bead:DNA) ratio. Recovered DNA was quantified using PicoGreen and reactions normalized to 5ng DNA (or entirety of the reaction if <5ng DNA was recovered) before preparing sequencing libraries (CUTANA CUT&RUN Library Prep kit; EpiCypher #14-1001). All autoCUT&RUN steps were optimized / performed on Tecan Freedom EVO robotics platforms with gentle rocking for incubation steps and magnetic capture for media exchange / washing steps. Sequencing was performed on an Illumina NextSeq2000 to generate 2x50bp paired-end reads.

### Immunofluorescence Staining

10T cells were seeded on 8-well chamber slides and after a PBS wash, fixed with paraformaldehyde (4% for 15 min) and permeabilized with Triton X-100 (0.5% for 15 min). After a PBS wash, slides were blocked with 3% BSA on rotator for 30 min and washed with PBS and incubated with anti-Flag (Sigma, #F1804) or anti-HA antibody (Biolegend, #901501) overnight at 4 °C (1:200 dilution in 3% BSA). Samples were washed three times with PBS before incubating with Alexa Fluor anti-mouse IgG secondary (ThermoFisher) for 1 h (1:1000 in 3% BSA). Samples were then counterstained with 4′,6-diamidino-2-phenylindole (DAPI) (ThermoFisher) solution in PBS for 15 min and washed three times with PBS. Slides were inverted onto gel mount on microscope slides, viewed and photographed with an Olympus BX43 microscope.

### Immunolabeling with fluorescent in situ hybridization (ImmunoFISH)

Cells were grown on glass #1.5 coverslips coated with Poly-D-lysine in 24 well plates. For imaging, cells were washed with PBS and fixed with 4% formaldehyde solution (10 min at RT). After washing twice with PBS, cells were permeabilized with 0.1% Triton X-100 in PBS (5 min at RT) and immunolabeled with anti-HA (Biolegend, 901501; 1:500 in PBS) for 1 hr at RT. After three washes with PBS, cells were incubated with secondary antibody (1 hr at RT), and again washed three times with PBS. Cells were re-fixed with 4% formaldehyde for 10 min and then hybridized with Stellaris RNA FISH probe for mouse Xist with Quasar 570 dye (BioSearch Tech, SMF-3011-1). First, cells were washed with PBS and incubated with Stellaris Wash Buffer A (SMF-WA1-60) for 5 min. Coverslips were then transferred to a dark, humidified container and incubated (37 °C for 4 hr) with Stellaris FISH hybridization buffer (SMF-HB1-10) containing the probe (1:100). Next, cells were washed with Wash buffer A (30 min at 37 °C), counterstained with DAPI in Wash Buffer A (30 min at 37 °C), and then incubated with Stellaris Wash Buffer B (SMF-WB1-20; 5 min at RT°) before being mounted onto a slide. Images were taken and deconvoluted using a DeltaVision Restoration Microscopy system (GE Healthcare) with an Inverted Olympus IX-70 microscope using a 40x objective.

### RNA-seq

Total RNA was isolated using TriZol and precipitated with isopropanol. Total RNA was quality checked using Bioanalyser (RIN>9) and libraries generated using NEBNext Ultra II Library Prep Kit (NEB, E7770) as per manufacturer’s protocol. Libraries were sequenced on an Illumina NextSeq 550 to generate 75-bp single-end reads.

### Organoid Culture Isolation and propagation

Murine organoids were generated as previously (49). Briefly, *K5Cre^ERT2^* transgenic mice were crossed to *R26tdTomato^lsl/lsl^* and *Trp53^loxP/loxP^* (Jackson Laboratory) to yield *K5Cre^ERT2^ ;R26tdTomato^lsl/lslt^ ;Trp53^loxP/loxP^*. Tamoxifen (Sigma-Aldrich; 0.25 mg/g body weight) was administered via oral gavage to 8-12 week-old littermates to knockout *Trp53* and activate TdTomato expression. Two weeks after tamoxifen treatment, 100 µg/ml of 4-Nitroquinoline *N*-oxide (4NQO; Sigma-Aldrich) in 2% propylene glycol (MP Biomedicals) was added to drinking water. Mice were exposed to 4NQO for 16 weeks and then followed up for 8-10 weeks to allow for tumorigenesis (48,49). Mice were euthanized and the esophagi harvested. A portion of the esophagus was reserved for histological evaluation of disease progression. The remaining tissue was placed in 5 U/µl dispase/PBS (BD Biosciences) for 10 minutes. The muscle layer was dissected from the epithelial layer and the latter treated with 0.25% trypsin and passed through a 40 µm cell strainer (BD Biosciences) to generate a single cell suspension. Using 24-well plates, 5000 cells were seeded per well in 50 μl of 3:1 Matrigel:organoid media solution. After solidification, 500 μl of organoid media (1:1 conditioned media prepared as described (77) and advanced DMEM media supplemented as in Table 2) were added per well and replenished every other day. To create transgenics, normal esophagus (EN) organoids were transduced with concentrated lentivirus to introduce DNMT3A1^WT^, DNMT3A1^W330R^ or DNMT3A1^W330R+R181E^, placed under G418 (300 μg/ml) after one week, and selected for two additional weeks. Methylation analysis was done after at least 21 days in culture. All experiments were performed under approved protocols from the Columbia University Institutional Animal Care and Use Committee (IACUC).

### Western Blot

Whole cell lysates were made in SDS Lysis Buffer (ThermoFisher) and resolved on 3-8% or 4-12% gradient SDS-PAGE gels (Invitrogen, NuPAGE). After transfer to a polyvinylidene difluoride (PVDF) membrane, same were blocked with 5% nonfat milk in PBST (PBS, 0.5% Tween-20) for 1h. Primary antibodies (1:1000) were added, incubated (overnight at 4 °C), washed in PBST, and detected with horseradish peroxidase-conjugated anti-rabbit IgG (1:10,000 in PBST). Antibodies used: DNMT3A (#3598, Cell Signaling), DNMT3L (#13451, Cell Signaling), β-Actin (#ab8227, Abcam), total-H3 (#ab1791, Abcam).

### Reduced Representation Bisulfite Sequencing (RRBS)

Genomic DNA (1ug) was collected from cells and sent to CD Genomics for RRBS sample preparation, library preparation and sequencing (Paired end 150bp).

### Enzymatic Methyl-Sequencing (EM-Seq)

EM-seq libraries were prepared from genomic DNA as previously (78) using the NEB Enzymatic Methyl-seq kit (New England BioLabs; P7120L). In brief, genomic DNA was purified using the NEB Monarch Genomic DNA Extraction kit (T3010L), and 10 ng sheared to an average fragment size of ∼300 bp (Diagenode Bioruptor Pico; B01060010) for enzymatic conversion / library preparation. Prior to sonication, fully 5mC methylated pUC19 (0.01 ng) and fully unmethylated lambda DNA (0.02 ng) were spiked into each reaction to monitor conversion efficiency. After nine cycles of PCR, library quality and quantification were assessed on the Agilent Tapestation 4200 using high sensitivity D1000 reagents (5067-5584 and 5067-5585). Libraries were sequenced (100M reads per biological replicate) on an Illumina NovaSeq 6000 to generate 100-bp paired-end reads.

### Fully defined nucleosomes

Fully defined (by mutation or PTM status) nucleosomes (*EpiCypher*) were assembled by salt-dialysis of semi-synthetic histones with 5’ biotinylated DNA (147bp of 601 nucleosome positioning sequence) (79) as previously (73,80) and confirmed by mass spectrometry and SDS-PAGE / immunoblotting (if an antibody was available). All Kub1 histones contain a native gamma-lysine isopeptide linkage (81,82).

### Nucleosome binding assay: dCypher™ Luminex

Biotinylated nucleosomes (Unmodified, Kub1, and acidic patch) were individually coupled to distinct magnetic avidin-coated xMAP bead regions (Luminex) at saturation and washed to remove excess. Individual [bead: Nuc] were adjusted to 1 million beads / mL, equimolar multiplexed, transferred to storage buffer (10 mM Cacodylate pH 7.5, 0.01% BSA, 0.01% Tween-20, 1 mM EDTA, 10 mM β-mercaptoethanol, 50% glycerol), and kept at -20 °C. Panel balance and individual [Nuc:bead] identity were confirmed in Luminex assays with anti-dsDNA (EMD Millipore #MAB030; 1/5, 1/50 and 1/500), anti-H3.1/2 (Active Motif #61629; 1/250, 1/1000 and 1/4000) and anti-PTM (EMD Millipore #04-263, CST #8240S and CST #5546S; all 1/250, 1/1000 and 1/4000). For the dCypher Luminex assay, multiplexed nucleosomes (1000 beads/well: the Targets) were used to examine the binding of serially diluted (two-fold: 1000 - 7.8 nM final) peptides (WT or mutant 6His-DNMT3A1 UDR [Biomatik]: the Queries). In brief, 50 µL of multiplexed dNuc panel was combined with 50 µL query peptide in UDR buffer (final: 20 mM Tris pH 7.5, 150 mM NaCl, 0.01% BSA, 0.01% NP40, 0.04 ug/mL salmon sperm DNA (salDNA; Invitrogen #15632-011), 1 mM DTT) in a black, flat-bottom 96-well plate (GreinerBio #655900), and incubated in the dark for one hr with shaking (800 rpm). Beads were captured / washed twice (100 µL of UDR buffer) on a plate-based magnet, 100 µL anti-6His (ThermoFisher #MA1-135; 1/1000) added and incubated in the dark for 30 min with shaking (800 rpm). Beads were captured / washed once (100 µL of UDR buffer) on a plate-based magnet, 100 µL of PE goat anti-mouse IgG (BioLegend #405307; 2 ug/mL) added and incubated in the dark for 30 min with shaking (800 rpm). After a final wash, beads were resuspended in 100 µL UDR buffer and Median Fluorescence Intensity (MFI) read on a FlexMap3D instrument (PerkinElmer), counting a minimum of 50-events per Target.

### DNMT3A1 N-terminal peptide

DNMT3A1 UDR peptides (76mers: aa159-228, N-terminal 6His tagged) were synthesized by Biomatik. Peptides were resuspended to 1 mg/ml in buffer containing 20mM Hepes pH 7.5, 0.5 mM EDTA and 1mM DTT.

### Preparation and reconstitution of Ubiquitinated nucleosomes

Plasmids containing wildtype histones were generous gifts from Dr. Karolin Luger, and mutant histone H2A-K119C was generated using the Q5 mutagenesis kit (NEB). Briefly, each histone was expressed in E. coli BL21(DE3) pLysS cells (Novagen, Cat# 69451), extracted from inclusion bodies, and purified sequentially by size exclusion and anion chromatography using previously published protocols (83). Purified histones were freeze-dried using a Sentry lyophilizer (VirTis).

The Ubiquitin plasmid DNA (pET-His-Ub G76C) was a generous gift from Dr. Tingting Yao. It was transformed into E. coli SoluBL21™ (Amsbio, Cat# C700200) competent cells and grown in 2xYT-Amp Media. Ubiquitin was expressed as soluble protein by inducing with 0.5 mM IPTG for 4 hours at 37°C upon the culture reaching OD600 = 0.4-0.6. Bacteria cells were harvested and lysed (AvestinEmulsiflexC3). Ubiquitin was purified on Ni-NTA agarose beads (Qiagen) Lysis Buffer (300 mM NaCl, 50 mM Tris pH 8.0, 10 mM Imidazole, 5 mM BME, 1x Protease Inhibitor) and Elution Buffer (300 mM NaCl, 50 mM Tris pH 8.0, 300 mM Imidazole, 5 mM BME) followed by HiTrap Q HP (GE Healthcare) liquid chromatography column with Buffer A (50 mM NaCl, 20 mM Tris pH 8.0, 0.2 mM EDTA, 10 mM BME) and Buffer B (1 M NaCl, 20 mM Tris pH 8.0, 0.2 mM EDTA, 10 mM BME). Purified ubiquitin was then dialyzed against water supplemented with 1 mM acetic acid followed by flash freezing in liquid nitrogen and lyophilized using a Sentry lyophilizer (VirTis).

Ubiquitinated H2A was generated following an established protocol (84). In brief lyophilized Ubiquitin (G76C) and histone H2AK119C were resuspended in Re-suspension buffer (10 mM Acetic acid, 7M Urea-Deionized) and mixed in the ratio of 2:1. Sodium Tetraborate, Urea, and TCEP were added to final concentrations of 50 mM, 6M and 5 mM. The mixture was incubated at RT° for 30 minutes. Then, an amount of crosslinker (1,3-dichloroacetone (Sigma, cat#167479) diluted in N,N’-dimethyl-formamide (Sigma, cat# D4551) equal to one-half molar ratio of total sulfhydryl groups was added to the solution and incubated on ice for 30 minutes. The reaction was stopped by the addition of BME to a final concentration of 5 mM. Then, the solution was diluted 10 times with denaturing binding buffer (Denaturing Binding Buffer: 50 mM Sodium phosphate (NaPi), 50 mM Tris pH 8.0, 300 mM NaCl, 6 M Urea, 10 mM Imidazole, 5 mM BME) and purified through Ni-NTA agarose beads (Qiagen) (Denaturing Elution Buffer: 50 mM NaPi, 50 mM Tris pH 8.0, 300 mM NaCl, 6 M Urea, 250 mM Imidazole, 5 mM BME). Purified Ubiquitinated-H2AK119C was dialyzed into water supplemented with 1 mM BME and lyophilized using Sentry lyophilizer (VirTis).

Nucleosomal DNA was generated from a plasmid containing eight copies of the Widom 601 positioning sequence, each flanked by an EcoRV site. This was transformed to *E. coli* DH5α (ThermoFisher) competent cells and grown in 2xYT-Amp media overnight. Cells were lysed and plasmid DNA purified using an established protocol (83). The 601 DNA fragment was excised using EcoRV (NEB) and fragments further separated from the vector backbone using repeated steps of PEG precipitation.

Nucleosome reconstitutions from individual histones were as described (83,85). Briefly, equimolar amounts of H3, H4, H2B, and ubiquinated-H2A were mixed and dialyzed into refolding buffer (2 M NaCl, 10mM Tris-HCl pH 7.5, 1 mM EDTA, 5 mM DTT). The octamer in refolding buffer was purified by size exclusion chromatography on a Superdex 200 16/600 column (GE healthcare). Nucleosomes were assembled by incubating purified Widom 601 DNA and histone octamers overnight with gradient salt dialysis using a peristaltic pump (Gilson Rapid Pump). Reconstitution was completed by dialyzing into Hepes buffer (20mM Hepes pH 7.5, 0.5 mM EDTA, 2mM DTT), and reconstitution efficiency was analyzed by 4.5% PAGE and quantified by DNA content.

### Cryo-EM sample preparation

A 5:1 complex between DNMT3A1 N-terminus peptide (25 µM) and H2A-Ub nucleosome (5 µM) was assembled in crosslinking buffer (50 mM HEPES, pH 7.9, 20 mM NaCl, 1 mM EDTA, 1 mM DTT), The complex was incubated for 2 hours at 4°C. The crosslinking reaction was performed by adding an equal volume of crosslinking buffer containing 0.06% glutaraldehyde and incubated for 1 hour at 4°C. The crosslinking reaction was quenched with 100 mM Tris pH 7.9 and dialyzed into buffer containing 20 mM Tris pH 7.9, 100 mM NaCl, 2 mM DTT, and concentrated for grid freezing. Cryo-EM grids of the peptide-nucleosome complex were prepared following the established protocol (86). Briefly, 3.0 μL of the samples with a concentration of 0.6 mg/mL was applied to Quantifoil gold grids (300 mesh, 1.2 μm hole size) glow-discharged for 25s. The grid was blotted for 3s with a blot force of 3 using Vitrobot Mark IV (FEI Company) at 4 °C and 100% humidity and plunge frozen in liquid ethane.

### Cryo-EM data acquisition, and processing

DNMT3A1 N-terminus peptide and H2A-Ub nucleosome complex data was collected on an FEI Titan Krios 300 kV equipped with a Gatan K3 Summit camera at a nominal magnification of 105,000x and a calibrated pixel size of 0.852 Å. Super-resolution images (0.426 Å/pixel) were collected as a movie stack for 2.4s, fractionated into 48 subframes, and the accumulated dose of 51.32 e-/Å2. A total of 3078 images were collected at a defocus range of 1.3 mm to 2.0 mm. The movies were motion-corrected, dose-weighed and binned to 0.852 Å/pixel using cryoSPARC’s ‘Patch Motion Correction’ function (87). The Contrast Transfer Function (CTF) was calculated using ‘Patch CTF Estimation (multi)’. Particles were picked using cryoSPARC’s ‘Blob Picker’ with a diameter between 80-120 Å, extracted from the micrographs in a 256 x 256 box and Fourier cropped to box 96 x 96 for the initial processing steps. Particles were subjected to 2D Classification into 50 classes, and the best particles were used to generate a model using the ‘Ab initio’ function, classified using ‘Heterogeneous Refinement’, and subsequently locally in 3D using ‘3D Classification’ with a wide mask focused on terminal DNA and Ubiquitin. The resultant map from these initial steps reflecting the best ubiquitinated-nucleosome features was refined, and its particles were re-extracted in a 256 x 256 box. These particles and the corresponding map were further locally classified with a focus on ubiquitin and refined to obtain the final reported map resolved at 2.88Å from 55,652 particles.

### Model building

The cryo-EM map of the DNMT3A1 N-terminus peptide and H2A-Ub nucleosome complex allowed for the unambiguous fitting of DNMT3A1 peptide, histones proteins, and DNA. Available X-ray structure of nucleosome from PDB:3TU4 was used for rigid-body fit into cryo-EM reconstructions for the nucleosome in Chimera (88), while the peptide was built de novo manually using Coot (89). The PDB was manually fit into the density and then locally optimized using UCSF Chimera’s "Fit in map" function (88). The complete model was refined using PHENIX (phenix.real_space_refine), using first only rigid body setting and then secondary structure, ADPs, rotamer, and Ramachandran restraints in 100 iterations (90). Ramachandran outliers and problematic regions were manually fixed in Coot. ChimeraX and PyMOL were used to prepare figures of the model and cryo-EM densities (89,91,92). Final model statistics are reported in Supplemental Data Table 3.

### Bioinformatic Analyses

ChIP–seq: Reads were aligned to mouse genome (GRCm38) using bowtie2 (93) (v2.4.4). Duplicated reads were removed by the *markdup* function from sambamba (94) (v0.6.8). DeepTools (95) (v3.5.1) was used for read depth normalization and visualization. Coverage tracks (bigwig) were generated by the bamCoverage function of DeepTools, normalizing to 1x depth (reads per genome coverage, RPGC) with bin size as 50bp and read extension as 200bp. Heatmap and enrichment plot of DNMT3A reads over CGIs were made with DeepTools with the *computeMatrix* (reference-point, -a 5000 -b 5000) and *plotHeatmap* functions. Genomic enrichment of DNMT3A reads were visualized using IGV (Integrative Genomics Viewer).

RNA-seq: Reads were trimmed using cutadapt (96) (v3.6) and pseudoaligned to the mouse genome (GRCm39) using kallisto (97) (v0.48.0, *quant*). Transcript-level abundances were read with tximport (v.1.28.0) to get gene-level abundances. Raw counts per gene were used with DEseq2 (98) (v.1.40) to find significantly differently expressed genes between samples (LFC >2 and q<0.01). For visualization with IGV, RNAseq reads were aligned to GRCm38 using hisat2 (v2.2.1) and converted to bigwig files with DeepTools function *bamCoverage*.

CUT&Tag and CUT&RUN data analysis: CUT&Tag and CUT&RUN reads were trimmed using cutadapt (v3.6) and mapped to the mouse genome (GRCm39 for CUT&Tag and GRCm38 for CUT&RUN) using bowtie2 (parameters --local --very-sensitive-local --no-unal --no-mixed --no-discordant --phred33 -I 10 -X 700) and duplicated reads were removed by the markdup function from sambamba. Coverage tracks (bigwig) were generated by the bamCoverage function of DeepTools with bin size as 50bp and normalizing to 1x depth (reads per genome coverage, RPGC). Heatmap and enrichment plot of H3K27me3, H2AK119Ub and H3K4me3 reads over promoters were made with DeepTools functions computeMatrix (reference-point, -a 5000 -b 5000) and plotHeatmap. Peaks of H3K27me3, H2AK119Ub and H3K4me3 CUT&Tag data were called using SEACR (99) (v1.3) and bedtools (100) (v2.26.0) intersect function was used to find promoters positive for each chromatin peak. Genomic enrichment of CUT&Tag and CUT&RUN signals were visualized using IGV.

### DNA methylation analysis (RRBS and EM-seq)

RBBS: Raw reads were filtered with trim galore (v0.6.6, --rrbs --paired for RRBS reads) and aligned to the mouse genome (GRCm39) using Bismark (101) (v0.23.1) with the functions *bismark* and *bismark_methylation_extractor* (--no_overlap --ignore 3 --ignore_r2 2 -- ignore_3prime 2 --ignore_3prime_r2 2). The resulting bismark coverage files were analyzed using methylkit (102) (v1.26) to calculate overall DNA methylation over all promoters and to find significantly differently methylated promoters (>10% methylation change and FDR<0.01, default Chi-square test). To compare methylation between RRBS samples, we restricted further analysis to CpGs that meet 10× coverage threshold across all samples and furthermore promoters (TSS +/- 500bp) with at least 5 CpG meeting the 10x coverage threshold. DeepTools functions *computeMatrix* and *plotHeatmap* were used for visualization of methylation across TSS and CGIs (+/- 5000bp) using bismark generated bedgraph files. For visualization of promoter methylation on Integrated Genomics Visualizer tracks methylKit was used to generate bedGraph files for promoters with at least 5 CpGs meeting the 10x coverage threshold.

EM-Seq: Raw reads were filtered with trim galore (v0.6.6) and aligned to the mouse genome (GRCm39) using Bismark v0.23.1 with the functions bismark and bismark_methylation_extractor (--no_overlap --ignore 3 --ignore_r2 2 --ignore_3prime 2 --ignore_3prime_r2 2). The resulting bismark coverage files were analyzed using methylkit (v1.26) All CpGs were retained and 10kb bins or promoters (TSS +/- 500bp) with at least 10 CpGs were retained for analysis.

## Data Availability Statement

RRBS, EM-seq, ChIP-seq, RNA-seq, CUT&RUN and CUT &Tag data reported in this paper have been deposited at GEO. Accession numbers can be found at GSE247019.

## Acknowledgement

We thank colleagues for the generous supply of plasmids and materials (Methods). We thank members of the Lu lab for critical reading of the manuscript. This study was supported by NIH grants (R01CA266978 to C.L. and K-J.A.; R44GM119893, R44DE029633, R44CA212733 and R44HG011875 to EpiCypher; R01DE031873 and R35GM138181 to C.L.; T32GM007739 and F30CA224971 to D.N.W). K-J.A. was also supported by the Mark Foundation for Cancer Research. The content is solely the responsibility of the authors and does not necessarily represent the official views of the National Institutes of Health. Research reported in this publication used the resources of the Columbia HICCC Flow Cytometry Shared Resource, Confocal and Specialized Microscopy Shared Resource and the Judith P. Sulzberger Columbia Genome Center, funded in part through the Office of the Director, NIH under awards S10OD020056 and NCI Cancer Center Support Grant P30CA013696. Schematic illustrations were created using BioRender.

## Author Contributions

Conceptualization, K.H.G., S.A-A., M-C.K., K-J.A. and C.L.; Experiments and analysis, K.H.G., S.A-A., S.L.G., J.L.M., V.U.S.K., A.R.H., O.A.A., H.F.T., K.N., C.P.L., D.N.M., B.J.V., D.N.W., I.K.P., V.S., R.L., X.X., N.L., S.F.; Writing, K.H.G., S.A-A., K-J.A. and C.L. with input from all authors; Supervision, H.N., M-C.K., K-J.A., C.L.; Funding Acquisition, M-C.K., K-J.A., C.L.

## Competing Interests

EpiCypher is a commercial developer and supplier of reagents (*e.g.*, PTM-defined semi-synthetic nucleosomes; dNucs) and platforms (dCypher^®^, CUTANA^TM^ CUT&RUN) used in this study. S.L.G., J.L.M., V.U.S.K., A.R.H., O.A.A., I.K.P., H.F.T., K.N., C.P.L., D.N.M., B.J.V. and M-C.K. are employed by (and own shares in) EpiCypher. M-C.K. is a board member of EpiCypher.

---

**Table S1.**
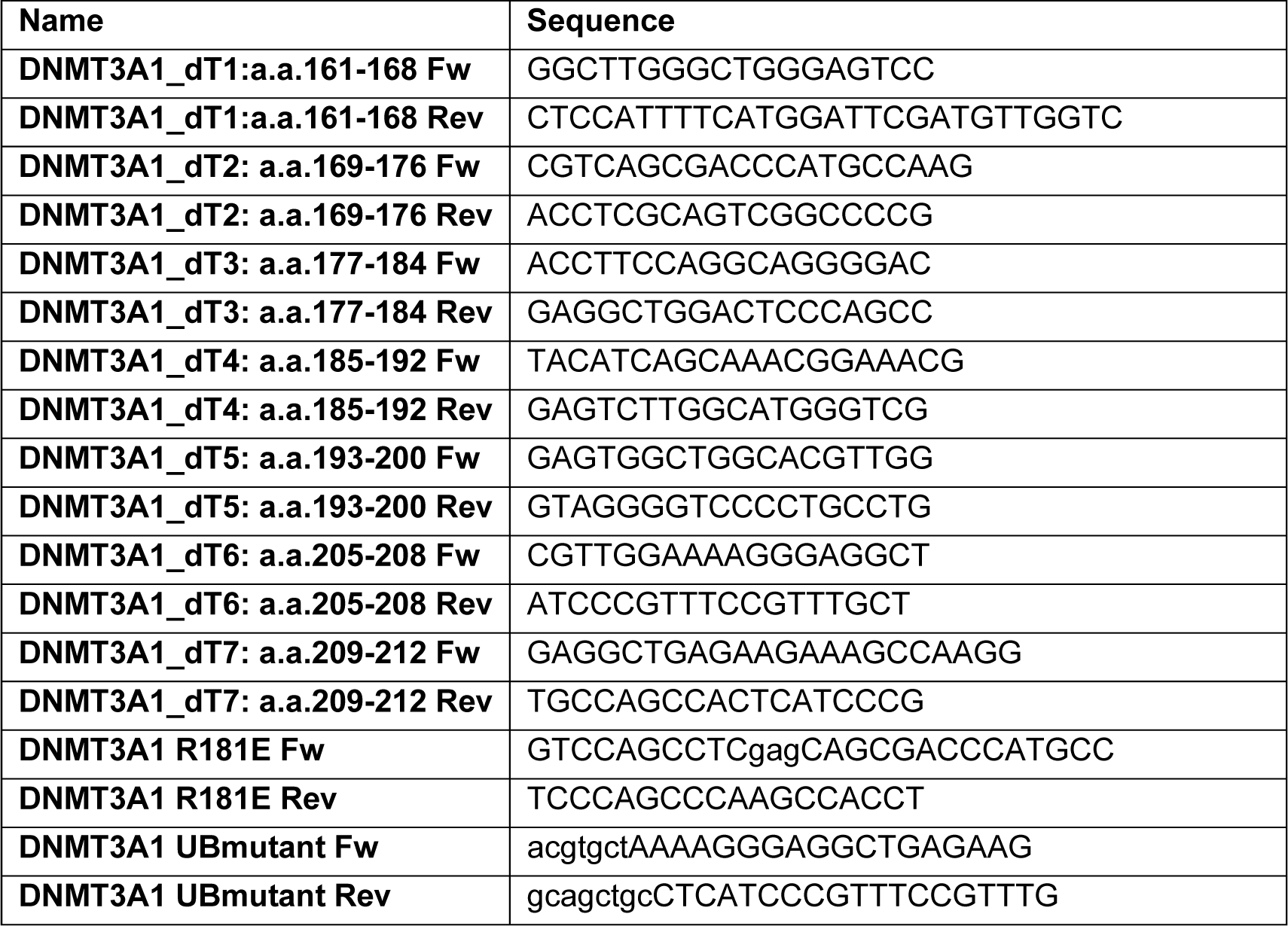
Primer Information.

**Table S2.**
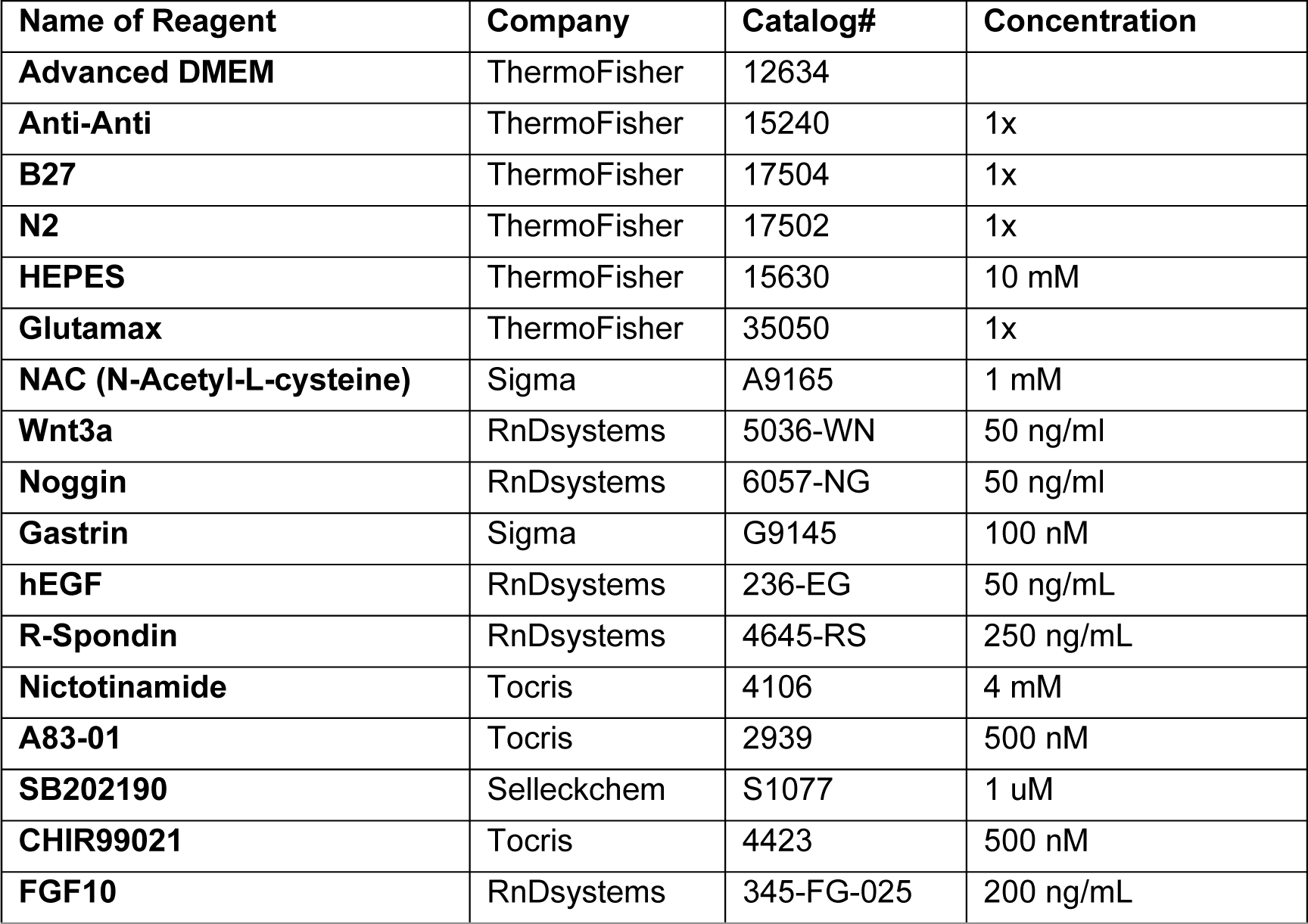
Organoid Media.

**Table S3.**
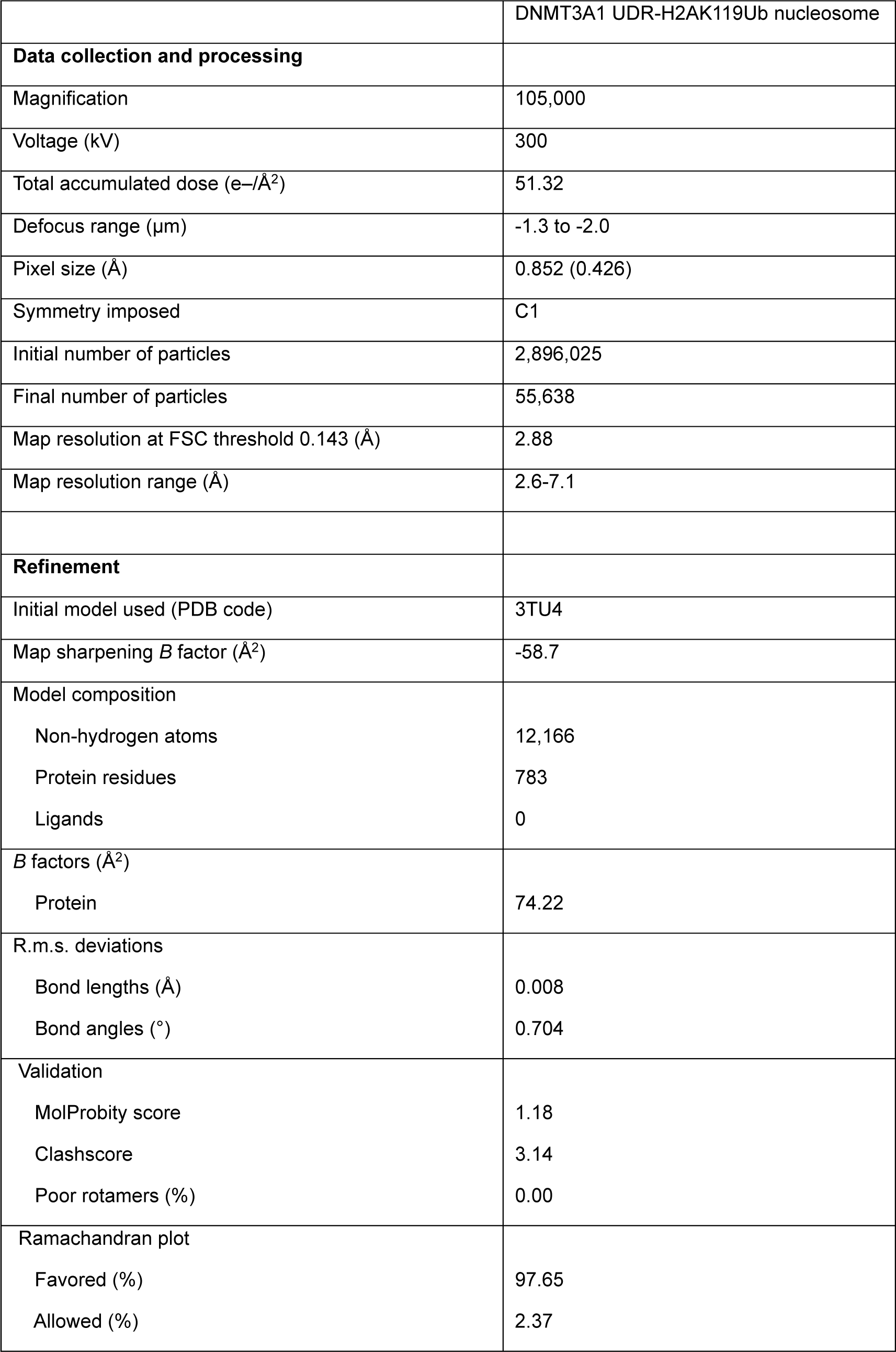
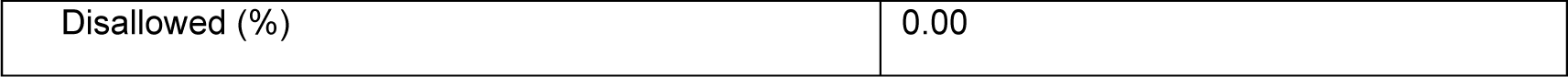
Cryo-EM data collection, refinement and validation statistics.

**Supplementary Fig. 1:**
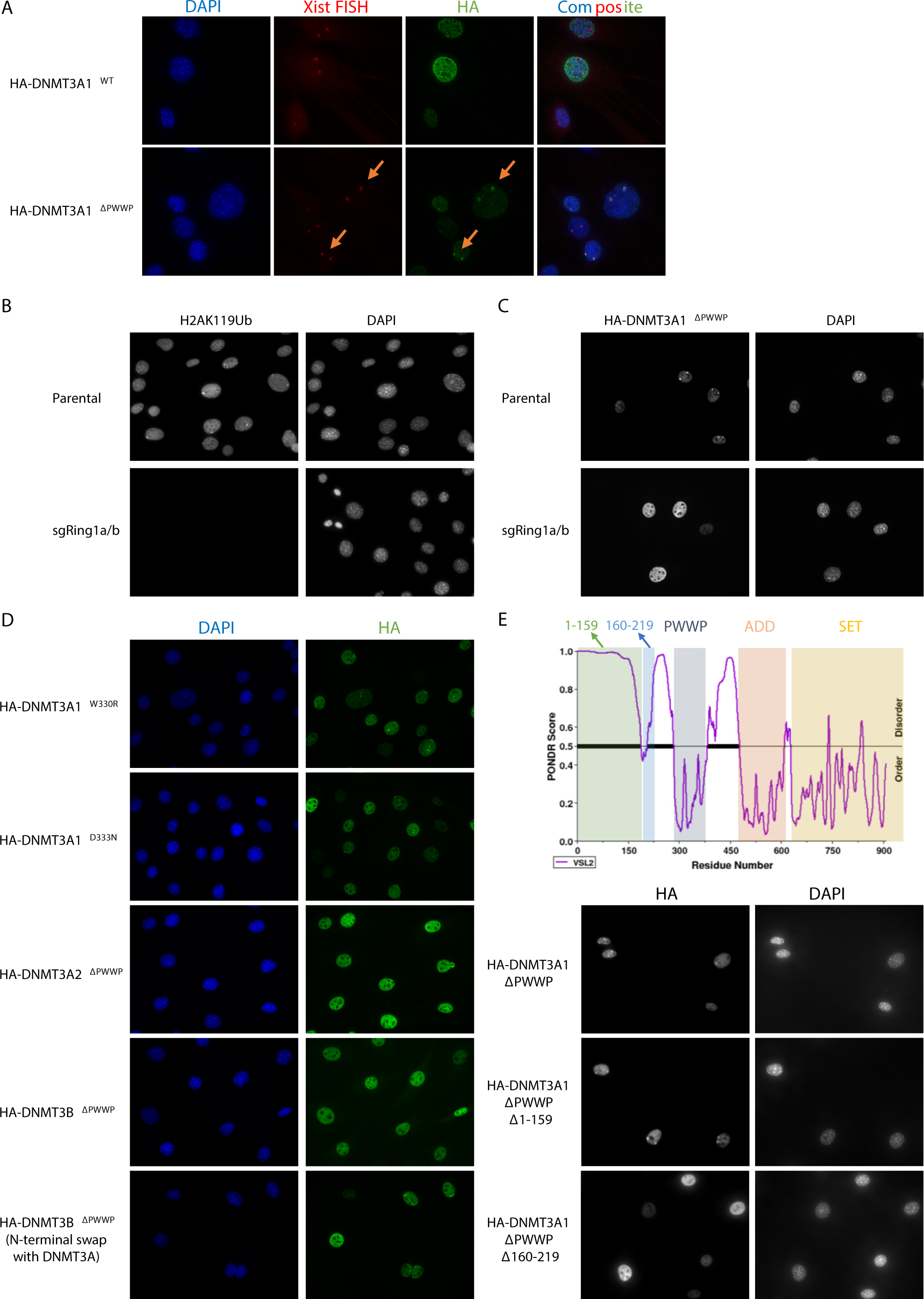
**Related to Figure 1. A)** Representative IF of DAPI, Xist RNA (FISH) and HA-DNMT3A1 in 10T cells expressing DNMT3A1^WT^ or DNMT3A1^ΔPWWP^ mutant. **B)** Representative IF of DAPI and H2AK119Ub in parental and sgRing1a/b 10T cells. **C)** Represen-tative IF of DAPI and HA-DNMT3A1 in parental and sgRing1a/b 10T cells expressing DNMT3A1^ΔPWWP^ mutant. **D)** Representative IF of DAPI and HA-DNMT3A1 in 10T cells expressing DNMT3A1^W330R^, DNMT3A1^D333N^, DNMT3A2^ΔPWWP^, DNMT3B^ΔPWWP^ or DNMT3B^ΔPWWP^ (N-terminal swap with DNMT3A) mutants. **E)** Representative IF of DAPI and HA-DNMT3A1 in 10T cells expressing DNMT3A1^ΔPWWP^, DNMT3A1^ΔPWWP+Δ1-159^ or DNMT3A1^ΔPWWP+Δ160-219^ mutants.

**Supplementary Fig. 2:**
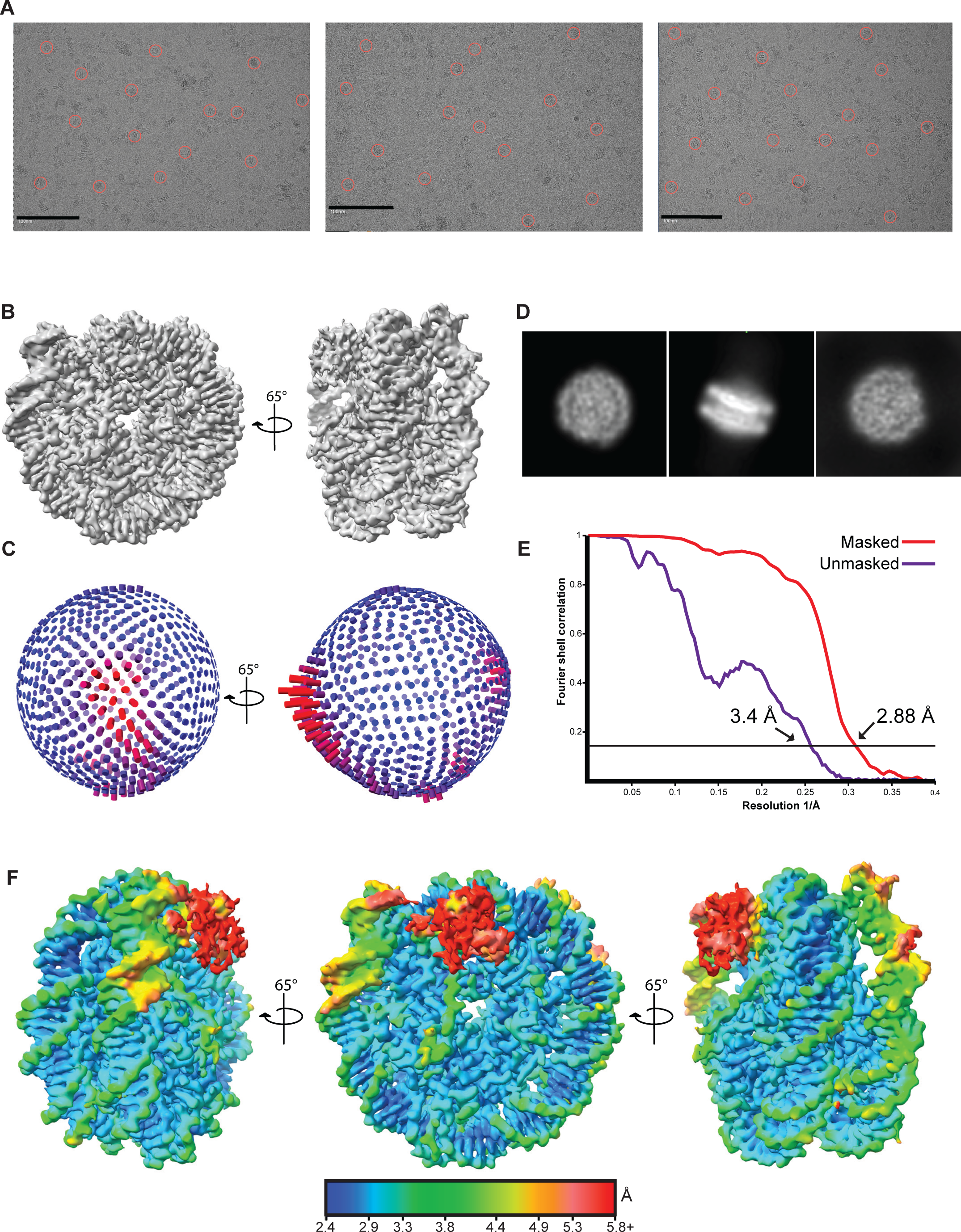
**Related to Figure 1. A)** Representative cryo-EM images from data collection with circles showing individual particles in the micrographs. **B)** 3D reconstruction of the N-terminal region of DNMT3A1 bound to H2AK119Ub nucleosome shown in two different orientations. **C)** Euler angle distribution of assignment of particles used to generate the final map with cylindrical length proportional to the number of particles assigned to a particular orientation. **D)** A representative 2D class averages selected from the dataset. **E)** Various views of the cryo-EM map colored according to local resolution (Å) measured at FSC of 0.143. **F)** Various views of the cryo-EM map colored by local resolution (Å).

**Fig s3.**
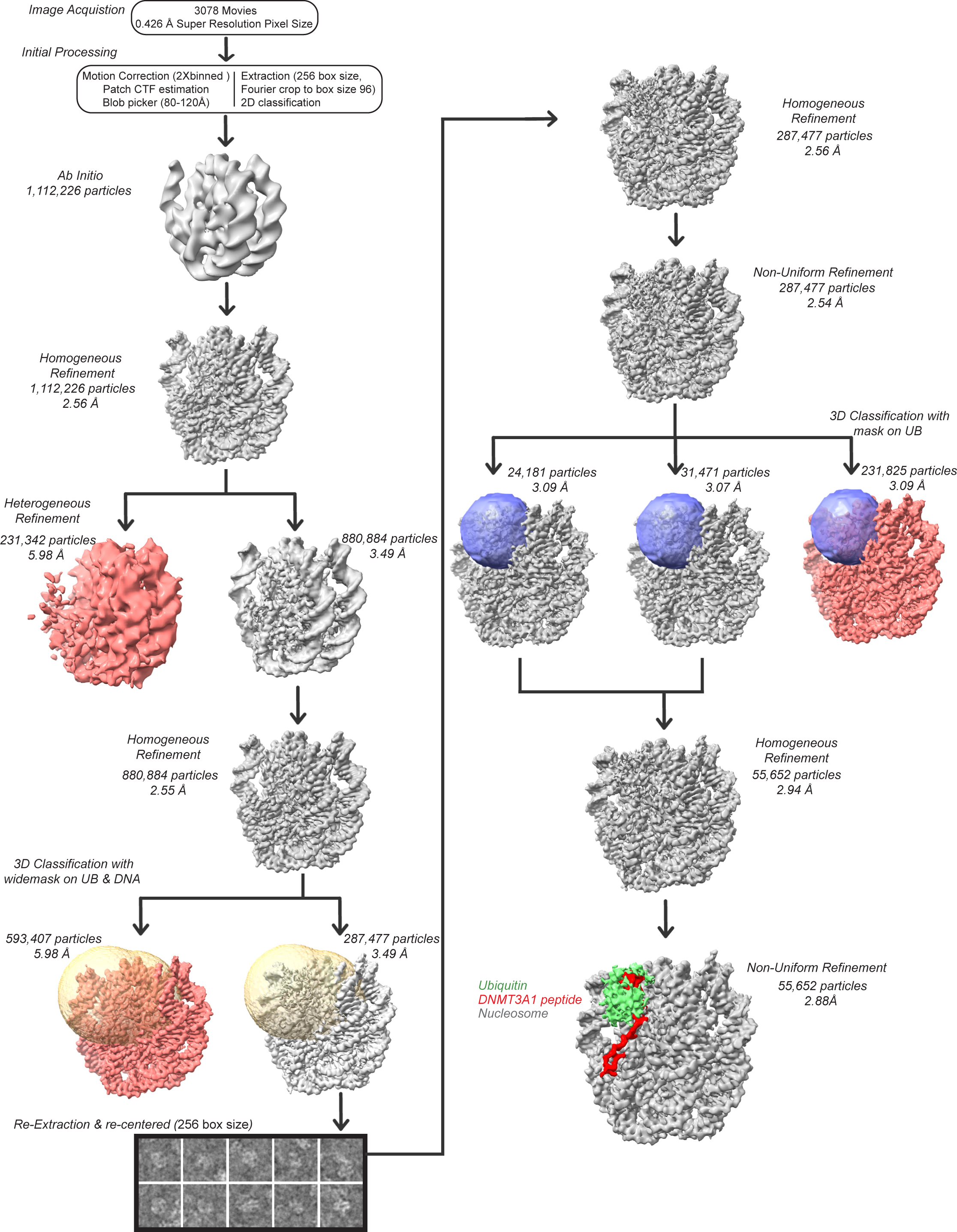
Data processing scheme of N-terminal region of DNMT3A1 bound to H2AK119Ub nucleosome.

**Supplementary Fig. 4:**
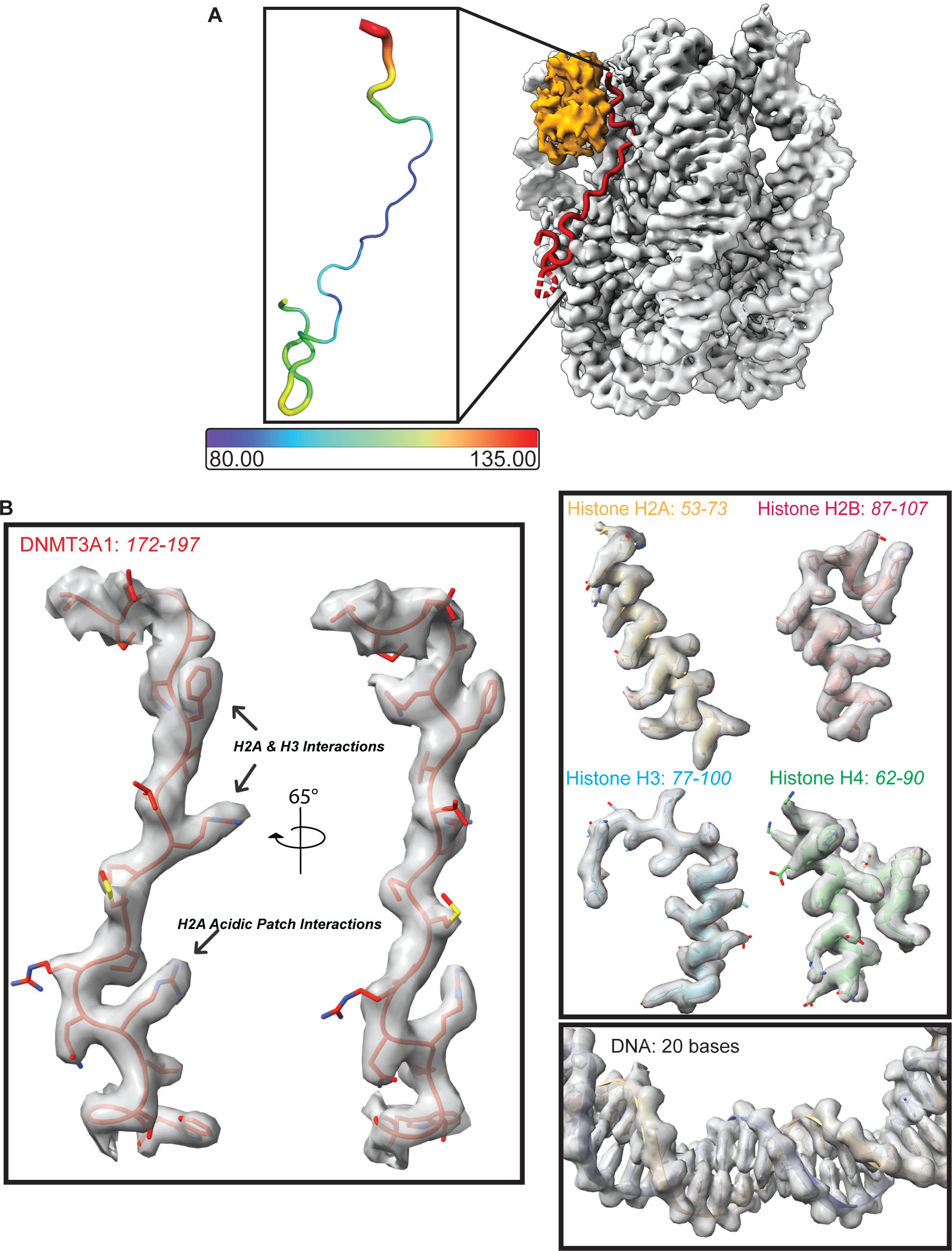
**Related to Figure 2. A)** Cartoon representation of the DNMT3A1 peptide from cryo-EM complex colored by residue B-factor. **B)** Cryo-EM densities and model fits of N-terminal region of DNMT3A1 peptide, and histones and DNA.

**Supplementary Fig. 5:**
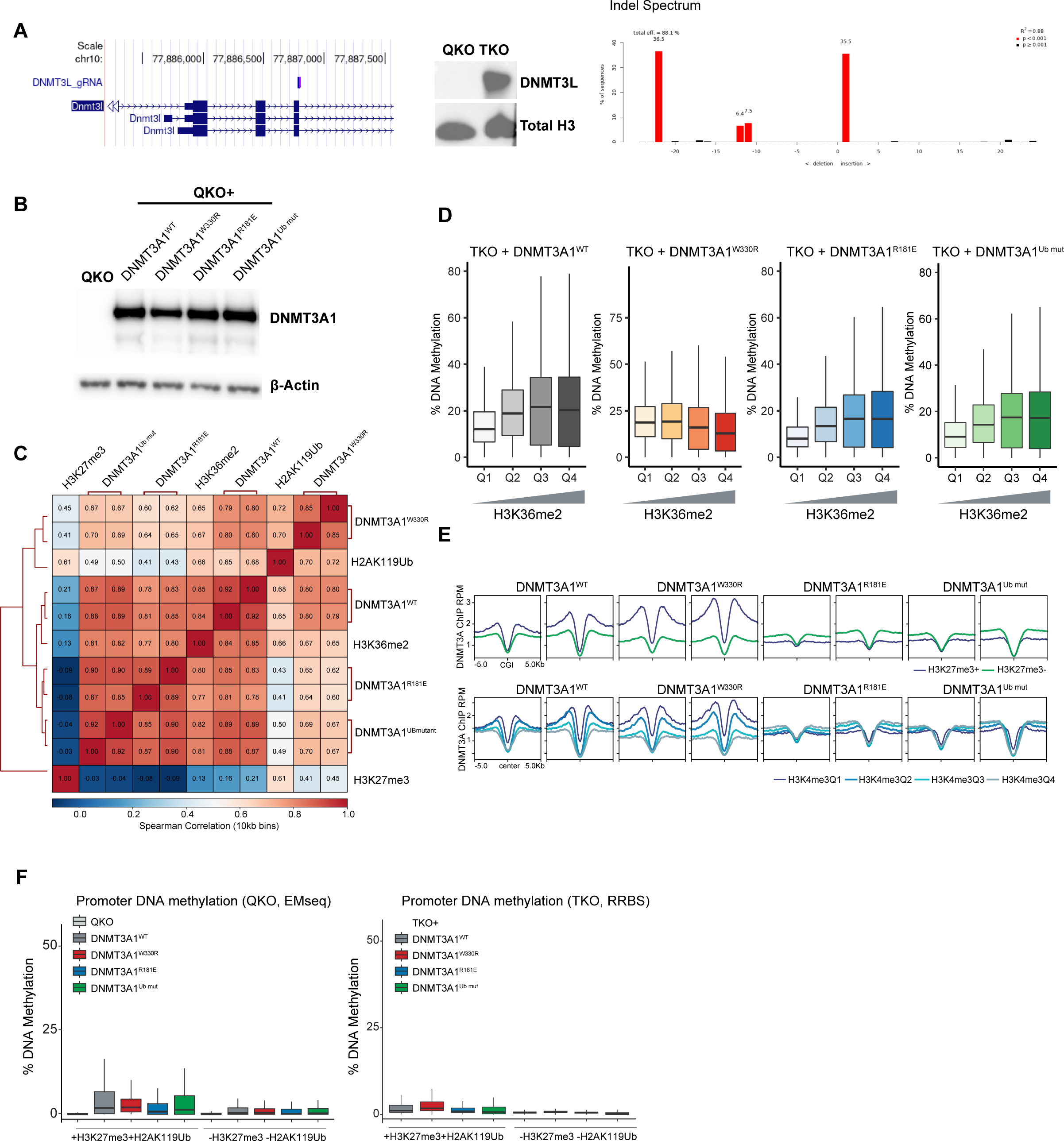
**Related to Figure 3. A)** Left: *Dnmt3l* locus showing the location of sgRNA. Middle: Western blot of DNMT3L in QKO and TKO mESCs with total H3 as loading control. Right: Validation of the *Dnmt3l* KO clone (QKO mESC) by TIDE (Tracking of Indels by Decomposition) analysis to quantify the frequency of small insertions or deletions. **B)** Western blot of DNMT3A in QKO mESCs and QKO mESCs expressing DNMT3A1^WT^, DNMT3A1^W330R^, DNMT3A1^W330R+R181E^ or DNMT3A1^W330R+Ub mut^. **C)** Correlation matrix of DNMT3A1 ChIP in QKO mESC expressing DNMT3A1^WT^, DNMT3A1^W330R^, DNMT3A1^W330R+R181E^ or DNMT3A1^W330R+Ub mut^, as well as H3K27me3, H2AK119Ub and H3K36me2 CUT&RUN in DNMT3A1^WT^ QKO mESC over 10kb bins. **D)** Percent DNA methylation measured by RRBS in DNMT3A1^WT^, DNMT3A1^W330R^, DNMT3A1^W330R+R181E^ or DNMT3A1^W330R+Ub mut^ TKO mESC per 10k bins grouped and ranked by H3K36me2 CUT&RUN enrichment (Q1 lowest to Q4 highest). **E)** Metagene plot showing enrichment of DNMT3A1 ChIP-seq reads in DNMT3A1^WT^, DNMT3A1^W330R^, DNMT3A1^W330R+R181E^ or DNMT3A1^W330R+Ub mut^ QKO mESC, centered at CGIs ± 5 kb and grouped by either H3K27me3 (top 20% vs. bottom 80%) or H3K4me3 (four quadrants). **F)** Left: Boxplot showing percent of promoter methylation in QKO mESC and QKO mESC expressing DNMT3A1^WT^, DNMT3A1^W330R^, DNMT3A1^W330R+R181E^ or DNMT3A1^W330R+Ub mut^ for both H3K27me3/H2AK119Ub-positive and -negative promoters. Right: Boxplot showing percent of promoter methylation in TKO mESC expressing DNMT3A1^WT^, DNMT3A1^W330R^, DNMT3A1^W330R+R181E^ or DNMT3A1^W330R+Ub mut^ for both H3K27me3/H2AK119Ub-positive and -negative promoters.

**Supplementary Fig. 6:**
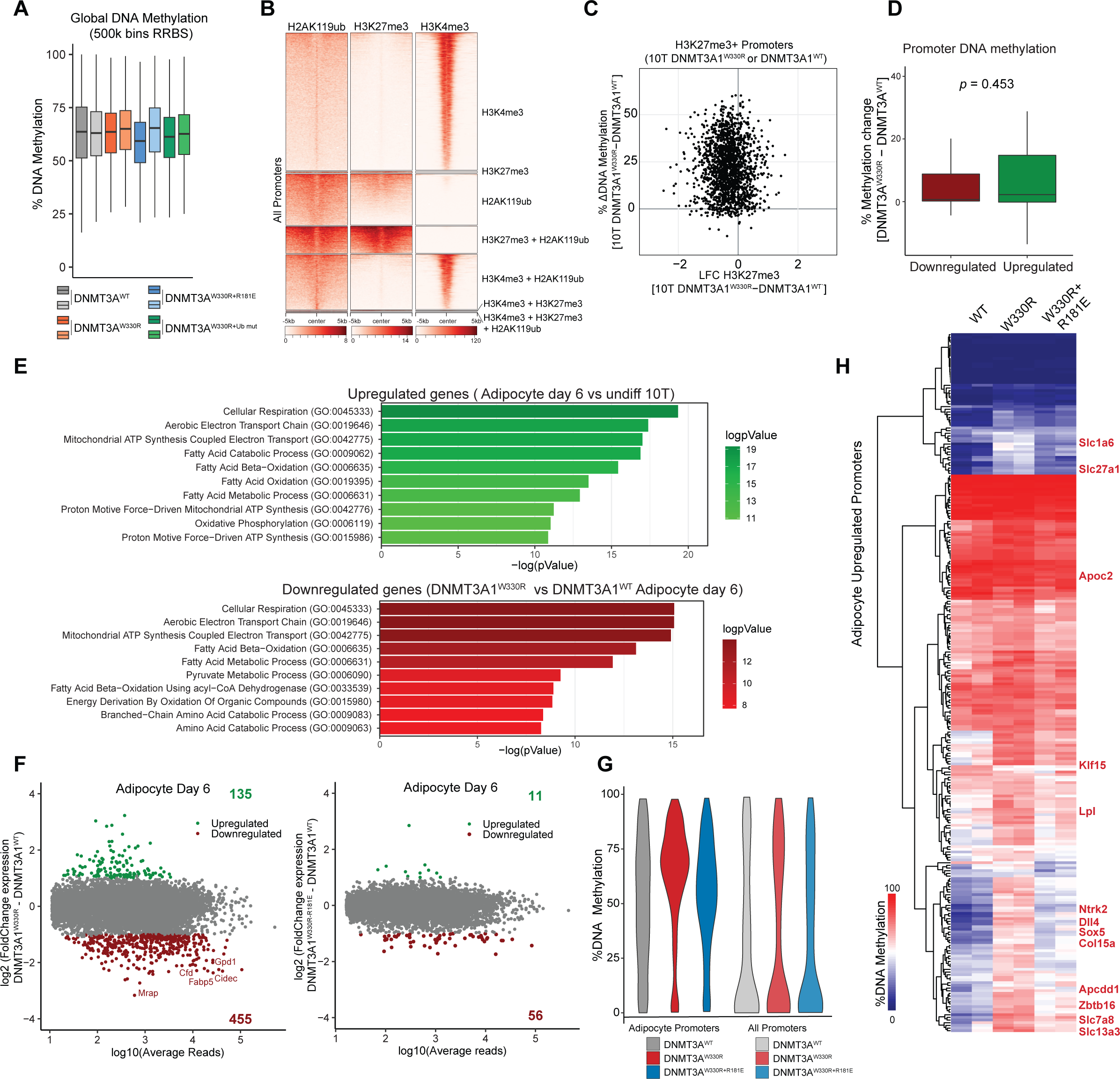
**Related to Figure 4 and 5. A)** Boxplot showing percent of global DNAme (500kb bins, measured by RRBS) in 10T cells expressing DNMT3A1^WT^, DNMT3A1^W330R^, DNMT3A1^W330R+R181E^ and DNMT3A1^W330R+Ub mut^. **B)** Heat map showing enrichment of H3K27me3, H2AK119Ub and H3K4me3 CUT&RUN reads at all promoters (TSS± 5kb) in 10T control cells. Promoters are classified into seven groups: H3K4me3^+^; H3K27me3^+^; H2AK119Ub^+^; H3K4me3^+^ and H2AK119Ub^+^; H3K27me3^+^ and H2AK119Ub^+^; H3K27me3^+^ and H3K4me3^+^; H3K4me3^+^ and H3K27me3^+^ and H2AK119Ub^+^ used in analysis for Fig 4C. **C)** Scatterplot comparing the changes in % DNAme (y-axis) and H3K27me3 (CUT&RUN, x-axis) between 10T cells expressing DNMT3A1^W330R^ and DNMT3A1^WT^ for all gene promoters positive for H3K27me3 in at least one condition. **D)** Boxplot comparing the differences in % promoter DNAme for both upregulated and downregulated genes between 10T cells expressing DNMT3A1^W330R^ and DNMT3A1^WT^. **E)** Barplot showing selected GO terms, ranked by -log(p-value) for, top: Upregulated genes in day 6 DNMT3A1^WT^ adipocytes compared to undifferentiated DNMT3A1^WT^ 10T cells (DEseq2, q>0.01 and LFC>2). Bottom: Downregulated genes in day 6 DNMT3A1^W330R^ vs. day 6 DNMT3A1^WT^ adipocytes (DEseq2, q>0.01 and LFC< -2). **F)** MA-plot showing log10 of average normalized RNA-seq reads for each gene (x-axis) and log2 fold change (LFC) of gene expression (y-axis) between (left) DNMT3A1^W330R^ and DNMT3A1^WT^ 10T cells on day 6 of adipocyte differentiation; (right) DNMT3A1^W330R+R181E^ and DNMT3A1^WT^ 10T cells on day 6 of adipocyte differentiation. Significantly differentially expressed genes (by DEseq2) are highlighted in green (LFC>2 and q<0.01) or red (LFC<-2 and q<0.01). **G)** Violin plots showing the distribution of % promoter DNAme (measured by RRBS) in 10T cells expressing DNMT3A1^WT^, DNMT3A1^W330R^, DNMT3A1^W330R+R181E^ for all promoters, or promoters of genes upregulated during adipocyte differentiation. **H)** Heatmap showing % promoter methylation (measured by RRBS) in 10T cells expressing DNMT3A1^WT^, DNMT3A1^W330R^, DNMT3A1^W330R+R181E^ for promoters of genes upregulated during adipocyte differentiation.

**Supplementary Fig. 7:**
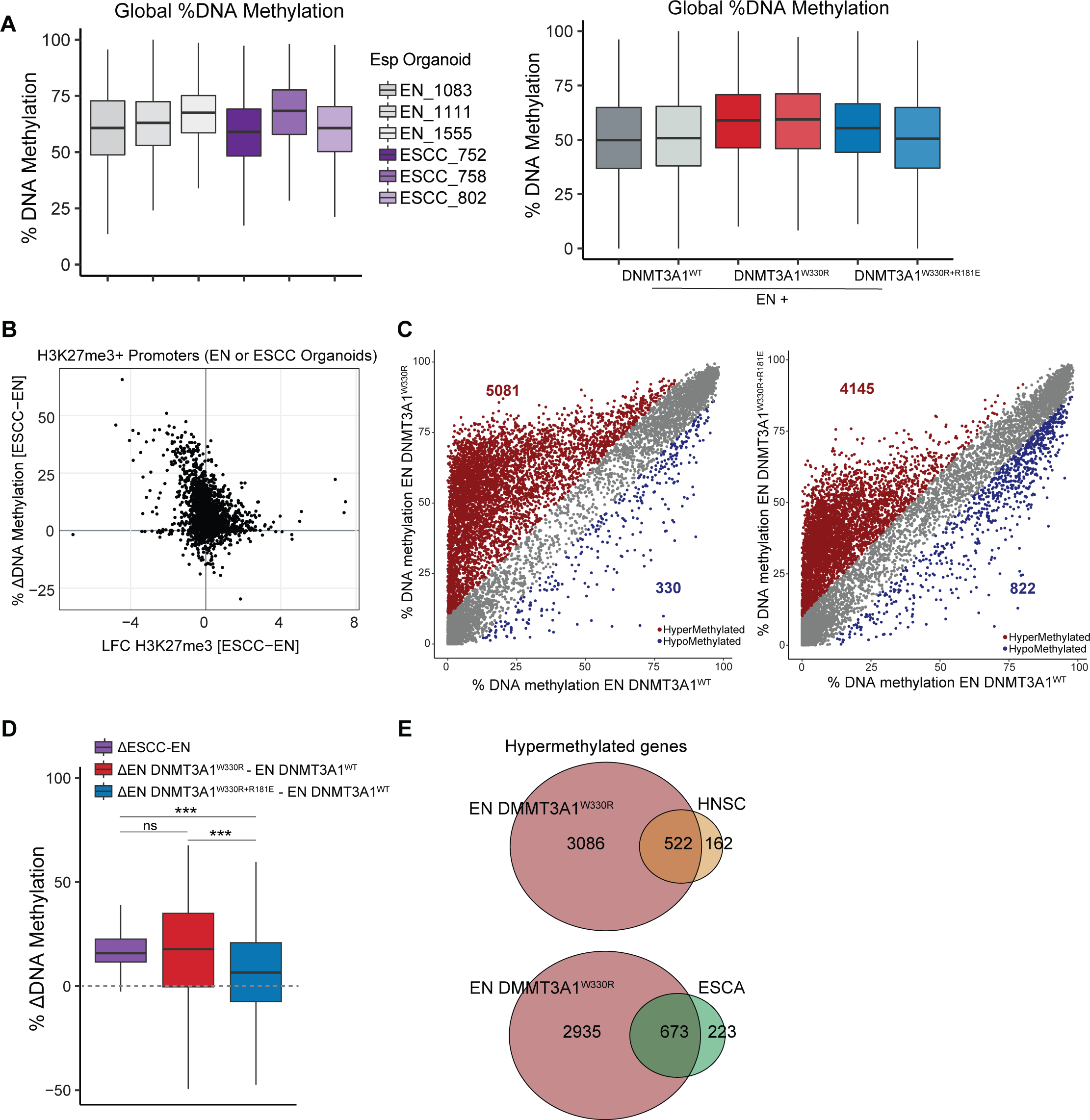
**Related to Figure 6. A)** Left: Boxplot showing percent of global DNAme (500kb bins, measured by RRBS) in 3x EN and 3x ESCC organoids. Right: Boxplot showing percent of global DNAme (500kb bins, measured by RRBS) in EN organoids expressing DNMT3A1^WT^, DNMT3A1^W330R^, DNMT3A1^W330R+R181E^. **B)** Scatterplot comparing the changes in % DNAme (y-axis) and H3K27me3 (CUT&RUN, x-axis) between EN and ESCC organoids for all gene promoters positive for H3K27me3 in at least one condition. **C)** Scatter plots showing % DNAme (measured by RRBS) at all promoters (TSS ± 500bp) in EN organoids expressing DNMT3A1^W330R^ or DNMT3A1^W330R+R181E^ versus EN organoids expressing DNMT3A1^WT^. Each dot represents a single promoter. Significantly hyper- (>10% and q<0.01) and hypo-methylated (>-10% and q<0.01) promoters are colored in red and blue, respectively. **D)** Boxplot showing the differences in % DNAme between ESCC and EN organoids (purple), EN organoids expressing DNMT3A1^W330R^ and DNMT3A1^WT^ (red), EN organoids expressing DNMT3A1^W330R+R181E^ and DNMT3A1^WT^ (blue), for all ESCC hypermethylated promoters. Student t test, ns p>0.05, * p<0.05, ** p<0.01, *** p<0.001. **E)** Venn diagrams showing the overlap between hypermethylated promoters in EN organoids expressing DNMT3A1^W330R^ and hypermethylated promoters in HNSC (top) or ESCA (bottom) patient samples.

